# The SF3B1 inhibitor pladienolide B massively inhibits DNA damage signaling and repair and counteracts resistance to platinum salts in Non-Small Cell Lung Cancer

**DOI:** 10.64898/2026.02.17.706284

**Authors:** Nour Jamal-El-Hussein, Shipra Chaudhary, Elodie Montaudon, Fariba Nemati, Aurélie Genoux, Hélène Polveche, Rania El Botti, Caroline Barette, Tao Jia, Didier Auboeuf, Didier Decaudin, Beatrice Eymin

## Abstract

**Background:** Lung cancer, including Non-Small Cell Lung Carcinoma (NSCLC), is the leading cause of cancer mortality worldwide. Platinum salts are the gold standard chemotherapy for NSCLC but many patients develop resistance leading to disease progression. Identifying new therapeutic strategies to counteract resistance is crucial. Pharmacological compounds targeting core components of the spliceosome machinery have emerged as promising anti-cancer agents. However, their mechanisms of action remain to be elucidated in NSCLC.

**Methods:** Various NSCLC cell lines were used in 2D and 3D cultures or clonogenic assays. NSCLC Patient-Derived Xenografts were also used. *SF3B1* was silenced by siRNA. Flow cytometry was performed to analyze cell cycle distribution and apoptosis. Western-blot, immunofluorescence, SIRF analysis and DNA repair assays were done to assess globally the DNA damage response. RNA-Seq, RT-qPCR and RT-PCR studies were performed to identify gene and splicing events impacted by SF3B1 inhibition. Publicly available transcriptomic and proteomic data were analyzed.

**Results:** SF3B1 is a core component of the spliceosome machinery. We show that NSCLC cells with acquired resistance to platinum salts are vulnerable to pladienolide B, a SF3B1 inhibitor, or *SF3B1* knock-down. Importantly, pladienolide B also slows down tumor growth of NSCLC Patient-Derived Xenografts (PDXs) poorly responsive to platinum salts. Mechanistically, we show that pladienolide B leads to genomic instability and apoptosis, that correlate with early transcription-dependent replication stress and DNA-PKcs activation, followed by the shutdown of ATR/DNA-PKcs-dependent signaling. In addition, pladienolide B profoundly regulates the expression and/or splicing, particularly exon skipping, of numerous genes involved in DNA repair, leading to decreased repair capacities of DNA double strand breaks. Although exon skipping events are mostly transient, skipping of exon 8 of *MLH3*, a gene involved in mismatch DNA repair, persisted along time. Finally, we show that pladienolide B counteracts resistance to platinum salts in NSCLC cells as well as PDXs, which correlates with enhanced *MLH3* exon 8 skipping and decrease of ATR, DNA-PKCs and MLH3 protein levels.

**Conclusions:** As a whole, our data highlight the targeting of SF3B1 as a potential therapeutic strategy, alone or in combination, in NSCLC patients who escape platinum salts-based chemotherapy.

## Introduction

Lung cancer remains the most commonly diagnosed malignancy and the leading cause of cancer-related mortality worldwide (1, 2). Non-Small Cell Lung Carcinoma (NSCLC) accounts for approximately 85% of all lung cancer cases (2). Although introduction of targeted therapies has led to considerable advances in therapeutic management of NSCLC patients with actionable mutations (3), chemotherapy continues to be a cornerstone treatment for unresectable and metastatic NSCLC patients who lack identifiable therapeutic targets (4). It is essentially based on the use of platinum salts (cisplatin, carboplatin) that form intra- and inter-strand adducts with DNA, and are thus potent inducers of cell cycle arrest and apoptosis (5). However, platinum salts suffer from considerable toxicity and many patients eventually relapse and become refractory to the drugs (6, 7).

Pre-mRNA splicing is a key post-transcriptional step in the regulation of gene expression (8). High-throughput RNA sequencing studies have now revealed that dysregulation of RNA splicing is a hallmark of cancer (9), with tumors harboring up to 30% more alternative splicing events than paired normal tissue from the same individual (10). Many causes leading to RNA splicing alterations in cancer have been identified, including heterozygous somatic mutations in splicing factors (11), abnormal expression of splicing regulatory proteins (12–14), or mutations in regulatory sequences in pre-mRNA (15). Further studies have shown that deregulated RNA splicing contributes to tumorigenesis by promoting proliferation, invasion, angiogenesis, immune escape, as well as resistance to therapies (16–18). All these observations have provided the rationale for the development of small molecule compounds that target core or regulatory components of the spliceosome machinery (19–21). However, despite encouraging pre-clinical results in various cancer types for some of these molecules, we clearly need to improve basic knowledge regarding their mechanisms of action, as well as to identify predictive biomarkers of response which could help to select patients who will better benefit from these therapies. In addition, as toxicity remains a major hurdle for their use in clinic, we need to identify drugs exhibiting additive/synergistic cytotoxic effects when combined with spliceosome modulators, a strategy which could allow to lower doses thereby to prevent adverse events.

The splicing factor 3B subunit 1 (SF3B1) is a core component of the spliceosome machinery that recognizes the branch site and the helix formed by base pairing between U2 small nuclear (sn)RNA and the pre-mRNA, facilitating branch point recognition and promoting the first step of the splicing reaction (22). SF3B1 mutations have been frequently identified in human cancer such as chronic lymphocytic leukemia, uveal melanoma, pancreatic ductal adenocarcinoma, or breast cancer (23, 24). Such mutations lead to cryptic 3′-splice site selection through the use of different branch points, which can activate oncogenic pathways or silence tumor suppressor genes (25). In addition, altered SF3B1 expression has been observed in several solid tumors, including breast, hepatocellular, prostate, and endometrial cancers (26). Natural compounds like spliceostatin A and pladienolide B, and their derivatives E7107 and H3B-8800, inhibit SF3B1 by binding to the interface between SF3B1 and PHF5A, another component of the U2snRNP (27, 28). Introns with weak branch points appear more sensitive to these compounds, including introns of genes related to cell division and apoptosis, which could support their anti-tumor effects (29–31). Consistently, all these drugs demonstrated anti-cancer effects in a wide range of cancer cells, including SF3B1-mutated cells (28, 32–34). Although E7107 and H3B-8800 both progressed to phase I/II clinical trials, E7107 was discontinued due to severe ocular side effects (35, 36) and H3B-8800 was halted due to insufficient efficacy. In preclinical studies, pladienolide B was shown to induce cell death in pancreatic ductal adenocarcinoma (PDAC) by increasing the expression of pro-apoptotic splice variants (e.g. BCL-X_S_), as well as to diminish PDAC-derived tumor growth in zebrafish and mice (34). Pladienolide B also inhibited tumor proliferation, migration, and colony formation in hepatocellular carcinoma (37), prostate cancer (33) or colorectal cancer (38). In contrast, the effects of pladienolide B or derivates in lung cancer cells remain poorly known.

In this study, we found that NSCLC cell lines with acquired resistance to platinum salts are vulnerable to pladienolide B-induced cell death. Mechanistically, we showed that pladienolide B activates an early transcription-dependent replicative stress response, which involves DNA-PKcs kinase activation, as well as widely impacts transcription and splicing, more particularly exon skipping, of numerous genes involved in DNA damage signaling and repair, including skipping of exon 8 of *MLH3*, a gene involved in mismatch repair. We further showed that pladienolide B counteracts resistance to cisplatin in NSCLC resistant cells lines which involves *MLH3* exon 8 skipping and correlates with a decrease of MLH3 protein level. More importantly, pladienolide B also slowed down tumor growth in NSCLC Patient-Derived Xenografts poorly responsive to platinum salts, with enhanced tumor growth inhibition when combined with platinum salts. In some PDXs, these effects of pladienolide B, alone or in combination, correlated with a decrease in ATR and/or DNA-PKcs protein level as well as enhanced skipping of *MLH3* exon 8. Altogether, these results suggest that targeting SF3B1 could provide therapeutic benefit in NSCLC patients with acquired resistance to platinum salts.

## Material and Methods

### Cells, Cell Culture and Reagents

Human Non-Small Cell Lung Carcinoma (NSCLC) cell lines (H460, A549, H1299, H2170, H3255, H460, PC9, CALU-1, H1975, H226, H358, HCC827) were authenticated by DNA STR profiling (ATCC cell line Authentification Service, LGC standards, Molsheim, France) and routinely tested for mycoplasma contamination. Parental and cisplatin-resistant cells derived from H460 and A549 NSCLC cells were obtained by continuous exposure to cisplatin according to (39). HCC827 and PC9 cells with acquired resistance to each generation of EGFR-TKIs (gefitinib, dacomitinib, osimertinib) were derived from parental sensitive EGFR-mutated PC9 or HCC827 cells following continuous exposure to increasing concentrations of each drug during 4-6 months as previously described (40) (41). Cells were cultured in RPMI-1640 medium/L-glutamax supplemented with 10% (v/v) Fetal Calf Serum (FCS) (42). Platinum salts-resistant H460 cells were continuously grown in presence of 20 µM carboplatin (cat#S1215, Selleckchem). In all experiments, carboplatin was removed before seeding. Pladienolide B (cat#sc-391691, Santa Cruz) and 5,6-dichlorobenzimidazole 1-ϕ3-D-ribofuranoside (DRB) (cat#D1916, Sigma Aldrich) were prepared at 10 mM in stock solution, stored at -80°C and dilute just before use in medium.

### 3D cultures

Spheroids were generated by seeding 1000 parental or resistant H460 cells**/**well in quadruplicate in complete medium into 96-well round bottom ultra-low attachment (ULA) spheroid microplates (Corning, Tewksbury, MA, USA). After 4 days, spheroids were treated or not with DMSO, pladienolide at 0.5 or 1 nM and let grown. Same treatments were repeated at day 11. Spheroid formation and growth at days 4, 7, 11 and 14 were assessed on an inverted microscope using MetaMorph software and multi-dimensional acquisition. Spheroid area in µm^2^ was calculated for each condition using Image J software.

### High-throughput screening of the effects of pladienolide B on cell viability

To assess the concentration-dependent effects of pladienolide B on cellular viability in 17 NSCLC cell lines, including platinum salts- and EGFR-TKIs- with secondary resistance models, we took advantage of the high-throughput screening platform for bioactive molecules (CMBA) at CEA Grenoble. 5x10^3^ cells/well were plated in 96-well plates and incubated overnight at 37°C before treatment with increasing concentrations of pladienolide B. After 72 hours, an automated viability assay was done based on fluorescence detection after incubation with PrestoBlue reagent (cat#A13262, Thermo Fisher).

### MTS assay

Cells (10x10^3^) were seeded in 96-well plates and allowed to adhere overnight at 37°C. Following 96 hours treatment of cells with increasing concentrations of cisplatin, MTS reagent [3-(4,5-Dimethylthiazol-2-yl)-2,5-diphenyltetrazolium bromide] was added to each well and incubated for 4 hours at 37°C. DMSO was added to each well and absorbance was recorded at 595 nm.

### Clonogenic survival assay

500 cells were seeded in 6-well plates and 24 hours later were exposed for ten to twelve days, to various drugs targeting distinct components of the DNA Damage Response signaling pathway at respective concentrations. These drugs included 10 µM olaparib, a PARP1/2 inhibitor (cat. #S1060, Selleckchem), 10 µM 6-OH-DOPA, a RAD52 inhibitor (cat. # H2380, Sigma Aldrich) or 5 µM BO2, a RAD51 inhibitor (cat. # SML0364, Sigma Aldrich) which was the only drug removed after 24 hours treatment. Twenty four hours after treatment, cells were treated with pladienolide B at 1 nM. Ten to twelve days later, the cells were washed with 1X PBS, fixed in 10% formalin for 5 min, washed two times in 1X PBS and stained with 1% methylene blue in 0.01 M sodium tetraborate decahydrate for 15 min at room temperature. Excess methylene blue was removed by washing with water and plates were dried overnight before colony counting. Colonies consisting of >50 cells were counted as positive. The number of colonies obtained in untreated condition was arbitrarily assigned to 100% survival.

### Active caspase-3 detection and cell cycle analysis by flow cytometry

Detection of active caspase-3 was performed by flow cytometry on an Accuri C6 flow cytometer (BD Biosciences) after cell fixation in Cytofix/Cytoperm™ and labelling with the phycoerythrin-conjugated monoclonal active caspase-3 antibody (BD Biosciences, Le Pont de Claix, France). For DNA content and cell cycle analysis, cells were fixed with 70% cold ethanol for 30 min on ice, treated with 20 µg/ml RNase A (cat. # R6513, Sigma Aldrich) for 20 min and stained with 10 µg/ml propidium iodide (cat. # P3566, Thermo Fisher Scientifics). Data from 10,000 cells were recovered and analyzed using the CellQuest software (BD Biosciences).

### Generation of stable clones for DNA repair analysis

Generation of stable clones for DNA repair analysis was done as previously described (43). Briefly, for the study of homologous recombination (HR), we used the pBL174-pDR-GFP plasmid (a kind gift from Dr Françoise Porteu, INSERM UMR1170, Villejuif). This plasmid contains two inactive genes coding for GFP under the control of a promoter. The 5’ gene is inactive because of the insertion of a cleavage site for I-SceI. The 3’ gene is inactive because it is deleted in both the 5’ and 3’ directions. When a DSB is produced by I-SceI (e.g. following transient transfection with the I-SceI-encoding pBL133 plasmid), recombination between these two inactive genes restores a functional GFP coding sequence. To establish stable clones, 10x10^6^ H1299 cells were transfected with 1 µg pBL174-pDR-GFP plasmid and clones were selected after 5 weeks culture in presence of 5 µg/ml puromycin. In order to select clones efficient for HR, stable clones were transiently transfected with pBL133-I-SceI plasmid encoding the I-SceI restriction enzyme for 4 days and detection of GFP-positive cells was done by flow cytometry. Four H1299 clones expressing GFP in 1.2 to 1.9% of cells as compared to cells transfected with control pcDNA3.1 plasmid were selected for further analyses. To assess the effects of pladienolide B, 0.2 x 10^6^ H1299-pBL174 cells were transfected with 0.5 µg pcDNA3.1 as a control or pBL133-I-SceI plasmid using XtremGene HP DNA transfection reagent (cat# 06366236001, Sigma Aldrich) according to the manufacturer’s instructions. After twenty four hours transfection, 5 nM pladienolide B or DMSO was added for 48 hours. Cells were fixed and the percentage of GFP-positive cells was quantified by flow cytometry on an Accuri C6 flow cytometer (BD Biosciences).

For the analysis of canonical-Non-Homologous End Joining (c-NHEJ), we used the pBL230 plasmid (a kind gift from Dr Françoise Porteu, INSERM UMR1170, Villejuif). This plasmid contains genes encoding the membrane antigens CD4 and CD8. CD8 is not expressed as it is in inverted orientation, and CD4 is not expressed because it is too far from the promoter. Two cleavage sites for I-SceI are present in non-coding sequences, which are in direct orientation generating cohesive ends between the two sites. When two DSBs are produced by I-SceI, rejoining of the DNA ends by exclusion or inversion leads to the expression of CD4 or CD8, respectively. 10x10^6^ A549 cells were transfected with 1 µg PBL230 plasmid and stable clones were selected after 6 weeks culture in presence of 10 µg/ml blasticidin. Clones having stably integrated PBL230 plasmid were first selected on the basis of the expression of H-2K^d^, a protein of class I major histocompatibility complex (MHC), using flow cytometry after cells fixation in 4% PFA in 1X PBS and staining using FITC anti-mouse H-2K^d^ antibody (clone SF1-1.1, cat#116605, Biolegend). An irrelevant FITC mouse IgG2a, k isotype was used as a negative control (clone MOPC-173, cat#400207, Biolegend). Three clones per cell line expressing H-2K^d^ in more than 80% cells were selected for further analyses. For detection of efficient c-NHEJ, 0.15x10^6^ A549-pBL230 clones were seeded in 6-well plates. Twenty four hours later, cells were transfected with 2.5 µg pBL133-encoding-I-SceI or pcDNA3.1 as a control using XTremGene HP (Sigma Aldrich) or Mirus Bio™ *Trans*IT™-LT1 (Fisher Scientific) transfection reagent according to the manufacturer’s instructions. After 48 hours or 72 hours transfection, 5 nM pladienolide B was added and cells were fixed in 2% PFA 96 hours after transfection. Detection of CD4 expression was done by flow cytometry, using a FITC anti-CD4 antibody (clone RM4-5; cat#553046, BD Biosciences). An irrelevant rat IgG2a, K isotype was used as a negative control (cat#553929, BD Biosciences).

### Premature Chromosomal Condensation (G2 PCC)

Chromatid breakage was analyzed in G2-phase cells using PCC to study the genomic instability consisting of chromosomal/chromatid breaks and exchanges. G2 PCC-like classical cytogenetics is a technique that allows the direct visualization and quantification of chromosomal aberrations. Exponentially growing H460 resistant cells were treated with 1 nM pladienolide B for 24 hours or 48 hours and allowed to grow at 37 ◦C. PCC was induced by incubating cells with 50 nM calyculin-A (cat#C-3987, LC laboratories) for 30 min before harvesting and processing. Calyculin A is a serine/threonine phosphatase inhibitor which efficiently induces Premature Chromosomal Condensations (PCCs) in interphase cells (44). For cytogenetic analysis, cells were incubated for 10 min at room temperature (RT) in a hypotonic solution, prepared by dissolving potassium chloride (Carl Roth GmbH & Co.) in water to obtain a 75 mM working solution, and fixed three times in Carnoy’s fixative. Carnoy’s fixative was prepared by mixing 3 parts methanol (Sigma Aldrich) and 1 part glacial acetic acid (Carl Roth GmbH & Co.) just before use. Fixed cells were dropped on clean glass slides and stained with Giemsa dye. Entellan (Merck) was used as a mounting medium. The analysis was carried out using bright field LSM510 META NLO confocal microscope (Zeiss, Oberkochen, Germany). Images were captured using AxioCam digital microscope camera. About 150 G2-PCCs were scored for excess PCC fragments at each time point and results from three independent experiments were combined. Breaks and gaps were scored, the latter when wider than the width of the chromosome.

### RNA Interference

Two sequences designed to specifically target human *SF3B1* RNA were purchased from Eurogentec (Angers, France) and were as follows: *Sf3b1≠1* 5’-GACAGCAGAUUUGCUGGAUACGUGA-3’; *Sf3b1≠2*5’-GCUUGGUCAGAAGAAGCCAGGAUAU-3’. For all RNA interference experiments, *mismatch* (control) siRNA used as a control was 5’-UCGGCUCUUACGCAUUCAA-3’. Cells were transfected with siRNA oligonucleotides duplex using RNAiMAX (Invitrogen, Cergy Pontoise, France) or JetPrime (Polyplus transfection, Ilkirch, France) reagent according to the manufacturer’s instructions. The cells were analyzed 72 hours post-transfection.

### Immunofluorescence studies

Immunofluorescence (IF) studies were performed on H460 parental or resistant growing cells. Briefly, 3x10^4^ cells were seeded onto 18-mm round coverslips pre-coated with 5 μg/ml fibronectin (cat#F2006, Sigma Aldrich), treated or not with 1 nM pladienolide B for 6 hours or 24 hours. Cells were then fixed with 4% PFA for 15 min at room temperature (RT), and permeabilized in 0.5% Triton X-100 for 20 min at RT. After three washes with 1X PBS for 5 min at RT, non-specific binding sites were blocked with 1X PBS, 10% bovine serum, and 0.3% Triton X-100 for 1 hour at RT before overnight incubation at 4°C with an anti-γH2AX antibody (cat# 05-636, Millipore) together with an anti phospho-53BP1(Ser1778) antibody (cat#2675, Cell Signaling Technology), or an anti RPA32 antibody (cat#52448, Santa-Cruz), overnight at 4°C. Next, the coverslips were washed three times with 1X PBS and 3% BSA, and a solution of 3% PBS–BSA with secondary antibody was added. Next, coverslips were placed in the dark.

After 2 hours, cells were rinsed three times with 1X PBS, and a drop of DAPI-Vectashield (Roche) was placed on a microscope slide to counterstain nuclei. A coverslip was placed on top of it and 24 hours later a multiphoton Zeiss (Oberkochen Germany) LSM510 META NLO confocal microscope was used to analyze IF experiments at 63X magnification. Images were acquired with AxioCam digital microscope camera and analyzed using ICY 1.7 software. Quantification of γH2AX, phospho-53BP1(Ser1778), and RPA32 foci number/size/intensity for each condition in each nucleus was performed using ICY or Image J software.

Immunofluorescence analyses of S-phase replicating cells were performed using click-iT EdU (ethynyldeoxyuridine) imaging kit (cat#C10340; Thermo Fisher Scientific) according to the manufacturer’s instructions. Briefly, 50 x 10^3^ platinum salts-resistant H460 cells were seeded onto 18 mm round coverslips pre-coated with 5 µg/mL fibronectin (cat#F2006, Sigma-Aldrich), treated or not for 6 hours with 5 nM pladienolide B and incubated with 10 µM EdU for 1 hour at 37°C. Cells were then fixed with 4% paraformaldehyde (PFA) for 15 min at RT, permeabilized in 0.5% Triton for 20 min at RT, before performing the Click-iT reaction using Alexa Fluor^TM^ 647-azide. Nuclei were counterstained using 6-diamidino-2-phenylindole (DAPI).

To analyze the role of active transcription in pladienolide B-induced γH2AX accumulation, 5x10^4^ H460R cells were seeded onto 18mm round coverslips pre-coated with 5 µg/mL fibronectin, treated or not for 6 hours with 5 nM pladienolide B in the presence or absence of 25 µM DRB. Cells were then fixed with 4% PFA for 15 min at RT, permeabilized in 0.5% Triton X-100 for 20 min at RT, and stained with anti-γH2AX antibody (clone JBW301, Millipore) for 1 hour at RT. Nuclei were counterstained using 6-diamidino-2-phenylindole (DAPI). Cells were observed using an Olympus microscope (x63 magnification). Images were captured with a Coolview CCD camera (Photonic Science) and digitally saved using Visilog software. Quantification of γH2AX foci number/size/intensity for each condition in each nucleus was performed using Image J software.

### SIRF analysis

SIRF for quantitative *“in situ”* analysis of protein interactions at DNA replication forks was performed in resistant H460 cells according to Roy et al (45). SIRF allows for the quantitative assessment of protein interactions with ongoing, stalled, and previously active replication forks using Proximity Ligation Assay (PLA) technology. Briefly, 4 x 10^4^ resistant H460 cells grown in log-phase were plated onto microscope chamber slides and treated or not with 5 nM pladienolide B for 6 hours. At the time of experiment, cells were incubated with 125 µM 5’-ethylene-2’-deoxyuridine (EdU) for 10 min, washed two times in ice-cold 1X PBS, permeabilized during 5 min at 4°C in CSK buffer (10 mM PIPES pH 6.8, 100 mM NaCl, 300 mM sucrose, 3 mM MgCl2, protease inhibitors cocktail) containing 0.5% Triton X-100 before fixation in 2% PFA for 15 min at room temperature. After fixation, PFA was discarded and the slides were extensively washed with 1X PBS before being incubated with the freshly prepared click reaction cocktail (2 mM copper sulfate, 9.5µM biotin-azide, 100 mM sodium ascorbate, 1 µM Alexa Fluor 488-azide in 1X PBS) for one hour at room temperature. EdU (cat#A10044), Alexa Fluor 488-azide (cat#A10266) and biotin azide (cat#B10184) were purchased from Thermo Fisher Scientific. After the click reaction, slides were blocked with blocking buffer containing 10% goat serum and 0.1% Triton X-100 in 1X PBS for 1 h at RT and incubated at 4°C overnight with the primary antibodies. The primary antibodies used in SIRF were: rabbit anti-biotin (cat#A150-109A; Bethyl Laboratory; 1/200e) or mouse anti-PCNA (cat#sc-56; Santa Cruz Technology; 1/200e). PLA was performed by incubating the slides for 1 h at 37°C in the presence of Duolink Rabbit minus and Duolink Mouse plus PLA probes (Sigma-Aldrich) diluted 1:5 in blocking solution. After washes, slides were incubated in ligation mix containing Duolink ligation stock (1:5) and ligase (1:40) at 37°C for 30 min. After washes, amplification was performed by incubating slides at 37°C for 100 min in an amplification mix containing Duolink amplification stock (1:5) and rolling circle polymerase (1:80). After extensive washes, slides were mounted in Prolong^TM^ Gold Antifade mountant with DAPI and analyzed using a multiphoton LSM510 META NLO confocal microscope (Zeiss, Oberkochen, Germany). Images were acquired with AxioCam digital microscope camera and analyzed using volocity software. All images are z-stacked.

### RIPA extracts for protein extraction and immunoblotting

For RadioImmunoPrecipitation Assay (RIPA) extracts, cells washed three times in 1X PBS were lysed in RIPA buffer [150 mM NaCl, 50 mM Tris–HCl (pH 8.0), 0.1% sodium dodecyl sulfate (SDS), 1% Nonidet P40, 0.5% sodium deoxycholate, 0.1 mM phenylmethanesulfonyl fluoride (PMSF), 2.5 μg/ml pepstatin, 10 μg/ml aprotinin, 5 μg/ml leupeptin, 0.2 mM Na_3_VO_4_] for 30 min on ice and sonicated at high frequency for 5 minutes (30 secs on 30 secs off cycle). Extracts were then pelleted by centrifugation at 13200 rpm for 30 min at 4°C. Immunoblotting experiments were performed as previously described (46). Antibodies used in this study are listed in Supplementary Table S1. Briefly, proteins were quantified and 20–40 μg proteins were separated by sodium dodecyl sulfate–polyacrylamide gel electrophoresis (SDS–PAGE) for 3 hours at 75 V using 3%-8% or 4%–12% precast polyacrylamide gels (Invitrogen) and 20X 3-morpholinopropane-1-sulfonic acid (MOPS) running buffer (Invitrogen). Next, proteins were transferred onto polyvinylidene difluoride (PVDF) or nitrocellulose membranes (Promega) for 16 hours at 35 V. The membrane was probed with primary antibodies overnight and washed three times with 0.1% PBS-Tween-20 followed by incubation with secondary antibodies for 2 hours. Membranes were scanned and analyzed using Fusion camera (Vilber). Quantification of the signals obtained for each protein was done using Image J software and the values were normalized according to the signal obtained for the loading control.

### RNA-Sequencing

Three biological replicate RNA samples from parental and resistant H460 cells treated or not with DMSO as a control or 3 nM pladienolide B for 8 hours were prepared using the NucleoSpin RNA isolation kit from Macherey-Nagel. RNA-Seq analysis was done by Novogen (Cambridge, United Kingdom). After RNA sample QC and directional library preparation (rRNA removal using Ribo-Zero kit), reads were generated on an Illumina NovaSeq 6000 platform (150 bp Paired End) allowing to obtain 15 G raw data per sample. Reads were mapped and filtered using TopHat2 (2.0.13), Samtools (0.1.19) and PrinSeq (0.20.4). Differential Gene Expression analysis was performed using DESeq2 (v1.26.0). Differences were considered as significant for absolute [log2 Fold Change] ≥ 0.4 and p ≤ 0.05. Differential splicing analysis was performed using replicate Multivariate Analysis of Transcript Splicing (rMATS) (47). The PSI value corresponding to exon inclusion rate was used to determine the sets of exons that were differentially spliced when comparing treated parental or resistant cells to untreated parental or resistant cells, respectively. Exons were defined as differentially regulated for absolute PSI variation (deltaPSI) ≥ 0.1, with *p*-values ≤ 0.05, as compared to the corresponding control. Exons were defined using FASTERDB (http://fasterdb.ens-lyon.fr/faster/home.pl). Gene Ontology Enrichment analyses were performed using the Gene Annotation and Analysis Resources Metascape (48).

### RNA extraction and RT-PCR analysis

Total RNA was extracted using NucleoSpin RNA kit (Macherey Nagel) according to the manufacturer’s instructions. 1 µg total RNA was reverse transcribed using iScript^TM^ Reverse Transcription Supermix (BioRad). The primers used for housekeeping gene were: GAPDH-Fw 5’-CGA-GAT-CCC-TCC-AAA-ATC-AA-3’; GAPDH-Rv 5’-ATC-CAC-AGT-CTT-CTG-GGT-GG-3. The list of all primers used in this study is presented in Supplementary Table S2. PCR conditions were 94°C, 2 min, 5 cycles (94°C, 30 s; 64°C, 30s; 72°C, 1 min), 5 cycles (94°C, 30 s; 61°C, 30s; 72°C, 1 min), 5 cycles (94°C, 30 s; 59°C, 30s; 72°C, 1 min), 25 cycles (94°C, 30 s; 56°C, 30s; 72°C, 1 min). Amplicons were detected on 1.8% agarose gels in 1X TBE.

### RT-qPCR analysis

One µg total RNA was reverse transcribed using iScript^TM^ Reverse Transcription Supermix (BioRad), diluted to 1:10 and 2 µl on 200 µl cDNA were used for qPCR. The primers were: GAPDH-Fw 5’-CGA-GAT-CCC-TCC-AAA-ATC-AA-3’; GAPDH-Rv 5’-ATC-CAC-AGT-CTT-CTG-GGT-GG-3’; SF3B1-Fw 5’-GTGGGCCTCGATTCTACAGG -3’ and SF3B1-Rv 5’-GATGTCACGTATCCAGCAAATCT-3’; PRKDC-Fw 5’-CTGTGCAACTTCACTAAGTCCA -3’ and PRKDC-Rv 5’-CAATCTGAGGACGAATTGCCT-3’; ATR-Fw 5’-ACCTCAGCAGTAATAGTGATGGA-3’ and ATR-Rv 5’-GGCCACTGTATTCAAGGGAAAT-3’; MLH3-Fw 5’-ACAAGCCAAATTGCGTTCTGG-3’ and MLH3-Rv 5’-TTCAGCATCAATACTGTTGAGGG-3’. qPCR was performed using SYBR Green Master Mix (BioRad) according to the manufacturer’s instructions on a BioRad CFX96 apparatus. Relative gene expression was calculated, for each sample, as the ratio of specific target gene to GAPDH gene (reference gene), thus normalizing the expression of target gene for sample to sample differences in RNA input.

### NSCLC Patient Derived Xenografts (PDXs) in *Nude* mice

These studies were performed in compliance with the recommendations of the French Ethical Committee and under the supervision of authorized investigators. The experimental protocol and animal housing followed institutional guidelines as put forth by the French Ethical Committee (Agreement number D-750602, France) and the ethics committee of the Institut Curie (Agreement number C75-05-18). Female *Nude* mice (Charles River, France) were xenografted with a tumor fragment of 20–40 mm3. Mice bearing tumors with a volume from 75 to 150 mm^3^ were individually identified and randomly assigned to the control or treatment groups. Eight NSCLC PDX models were used in this study: 2 squamous carcinomas (LCF2, LCIM1), 4 adenocarcinomas (ML5LC66, LCIM13, LCF15, LCIM10) and 2 large cell carcinomas (LCF26, ML1LC2). For ethical reasons, experiments were performed according to a related-“single-mouse” schedule in which at least three mice/PDX model bearing growing tumor were included per group (49, 50). Mice received either vehicle (control), cisplatin (4 mg/kg, ip, 1 x/3 weeks) and/or pladienolide B (2.5 mg/kg or 5 mg/kg, ip, D1-D4-D8-D11). Individual tumor volume and relative tumor volume (RTV) were calculated according to standard methodology (51). Briefly, tumor growth was evaluated with a caliper by measurement twice a week of two perpendicular diameters of tumors (a and b). Then, individual tumor volumes were calculated as: a x b^2^ / 2 in mm^3^, where a is the largest diameter and b is perpendicular to a. RTV was calculated as the ratio of the volume at time t divided by the initial volume at day 0 and multiplied by 100 [RTVt=(Vt/V0)x100]. Moreover, to evaluate the overall response rate (ORR) to treatments observed in all treated models according to individual mouse variability, we considered each mouse as one tumor-bearing entity. ORR was determined as the ratio RTVt/RTVc-1. A tumor was considered to be responding to treatment if ORR was below −0.5. Finally, to clearly evaluate the impact of treatments on the tumor progression, we evaluated the probability of progression by calculating doubling time and time for RTV x 4. At the time of first ethical sacrifice (when tumors reached 2000 mm^3^), frozen and formol-fixed tumor tissues were collected in all control and treated groups. Total RNA was extracted and 1 µg was reverse-transcribed to analyze *MLH3*-Exon 8 skipping by RT-PCR. RIPA whole protein extracts were also done and immunoblots were performed to detect ATR, P-ATR(Thr1989) or DNA-PKcs protein. Statistical significance of observed differences between the individual RTVs or ORR corresponding to the treated mice and control groups were calculated using the two tailed Mann–Whitney test. Median times to reach RTV=2 or RTV=4 were estimated on the basis of the Kaplan-Meier event-free survival distribution. The log-rank exact test, as implemented with GraphPad Prism, was used to compare the probability of progression between treatment and control groups.

### Data mining of publicly available databases

Protein expression levels of DNA-PKcs/PRKDC (UniProt Accession: P78527), ATR (UniProt Accession: Q13535), and SF3B1 (UniProt Accession: O75533) in lung cancer cell lines were retrieved from the Cancer Cell Line Encyclopedia (CCLE) proteomics dataset (https://gygi.hms.harvard.edu/publications/ccle.html). The data were downloaded as normalized protein quantification values. Spearman’s correlation coefficients were calculated to assess the relationship between protein expression levels of the indicated targets across lung cancer cell lines. Statistical significance was evaluated by GraphPad Prism 8.

Gene expression levels of *SF3B1* with *PRKDC* or exon usage values for *MLH3* exon 5 (ENSE00003488554) and exon 8 (ENSE00003584609, annotated exon 7 in ENSEMBL), were obtained from the publicly available TCGA Lung Adenocarcinoma (LUAD) dataset via the TCGA Splicing Variants Database (TSVdb) (http://www.tsvdb.com/index.html). Gene expression was expressed as log2-transformed RSEM-normalized read counts. Exon usage was calculated as the ratio of exon expression to total gene expression. Spearman’s correlation analysis was performed to assess associations between gene expression levels and exon usage. Statistical analysis was conducted using GraphPad Prism 8.

### Statistical analyses

Analyses were performed using the GraphPad Prism software (GraphPad Software Inc., San Diego, CA, USA) and data are presented as mean ± SD. Statistical comparisons between two groups or more were conducted with Mann-Whitney, paired or unpaired t test as mentioned in the text.

## Results

### Pladienolide B inhibits cell viability in NSCLC cells with acquired resistance to platinum salts, as well as tumor growth in NSCLC Patient-Derived Xenografts (PDXs) with primary resistance to platinum salts

We initially used the high-throughput screening platform for bioactive molecules at CEA Grenoble, and an automated PrestoBlue-based 72 hours cell viability assay, to screen the cytotoxic effects of pladienolide B in 11 NSCLC cell lines (A549, H1299, H2170, H3255, H460, PC9, CALU-1, H1975, H226, H358, HCC827), 2 NSCLC cell lines (derived from PC9 or HCC827) with acquired resistance to 3 distinct EGFR-TKIs (gefitinib, dacomitinib, osimertinib) and 2 NSCLC cell lines with acquired resistance to platinum salts (derived from H460 or A549, hereafter denoted H460R or A549R respectively). Resistant cellular models were obtained after chronic exposure to increasing concentrations of each EGFR-TKI or platinum salts for 4-6 months as described in the material and methods section. NSCLC cell lines exhibited different response to pladienolide B used at nanomolecular concentrations with some being less sensitive (e.g. A549, H3255) compared to others (e.g. CALU-1, H358) (Fig S1a). Interestingly, H460R or A549R cells, which were both resistant to cisplatin (Fig S1b), appeared more sensitive to pladienolide B as compared to parental cells (Fig 1a). Conversely, pladienolide B did not demonstrate distinct cytotoxic effects in PC9 or HCC827 cells with acquired resistance to either gefitinib, dacomitinib or osimertinib compared to parental PC9 or HCC827 cells (Fig S1c). Quantification of apoptosis further confirmed earlier and increased apoptosis in H460R or A549R cells treated with pladienolide B, as compared to parental cells (Fig 1b). Increased cell growth inhibition in response to pladienolide B in H460R cells was confirmed in clonogenic assay (Fig 1c), as well as in 3D spheroids culture (Fig 1d). As a whole, these results demonstrated that NSCLC cells with acquired resistance to platinum salts are vulnerable to pladienolide B-induced cytotoxicity. SF3B1 is the target of pladienolide B. SF3B1 was not differentially expressed between parental and platinum-salts resistant cells (Fig 1e). To eliminate off-target effects of pladienolide B, we silenced *SF3B1* by using two distinct siRNAs (Fig 1f, left panels). SF3B1 knock-down enhanced apoptosis in H460R or A549R cells as compared to H460 and A549 parental cells (Fig 1f, right panels). Finally, we tested the effects of pladienolide B on the tumor growth of 8 NSCLC Patients-Derived Xenografts (PDXs). These PDXs displayed distinct mutational status and were selected based on their histological subtype and their poor primary response to platinum salts (Table 1). Pladienolide B did not exhibit general toxicity *in vivo* based on the measurements of mice body weight (Fig S2a). Thirty percent of all PDXs displayed a moderate overall rate response (ORR< -0.5) to pladienolide B-induced tumor growth inhibition in a concentration-dependent manner (Fig 1g). According to ORR or the mean Relative Tumor Volume (RTV), LCF26, ML1LC2, LCIM1, LCIM10 and LCIM13 responded to pladienolide B with LCIM10 being the most responsive, while this was not the case for LCF2, LCF15 or ML5LC66 (Fig S2b-c). When probability of progression (RTV=2 or RTV=4) was calculated taking into account all PDXs (excepted LCIM1, n=7), we showed that pladienolide B slows down tumor growth in a dose-dependent manner (Fig 1h and Fig S2d). To our knowledge, this is the first demonstration of anti-tumoral effects of pladienolide B in NSCLC-derived PDXs. As a whole, these results demonstrated that NSCLC cells with acquired resistance to platinum salts and NSCLC PDXs that poorly respond to cisplatin are vulnerable to pladienolide B-induced cell or tumor growth inhibition.

**Figure 1.**
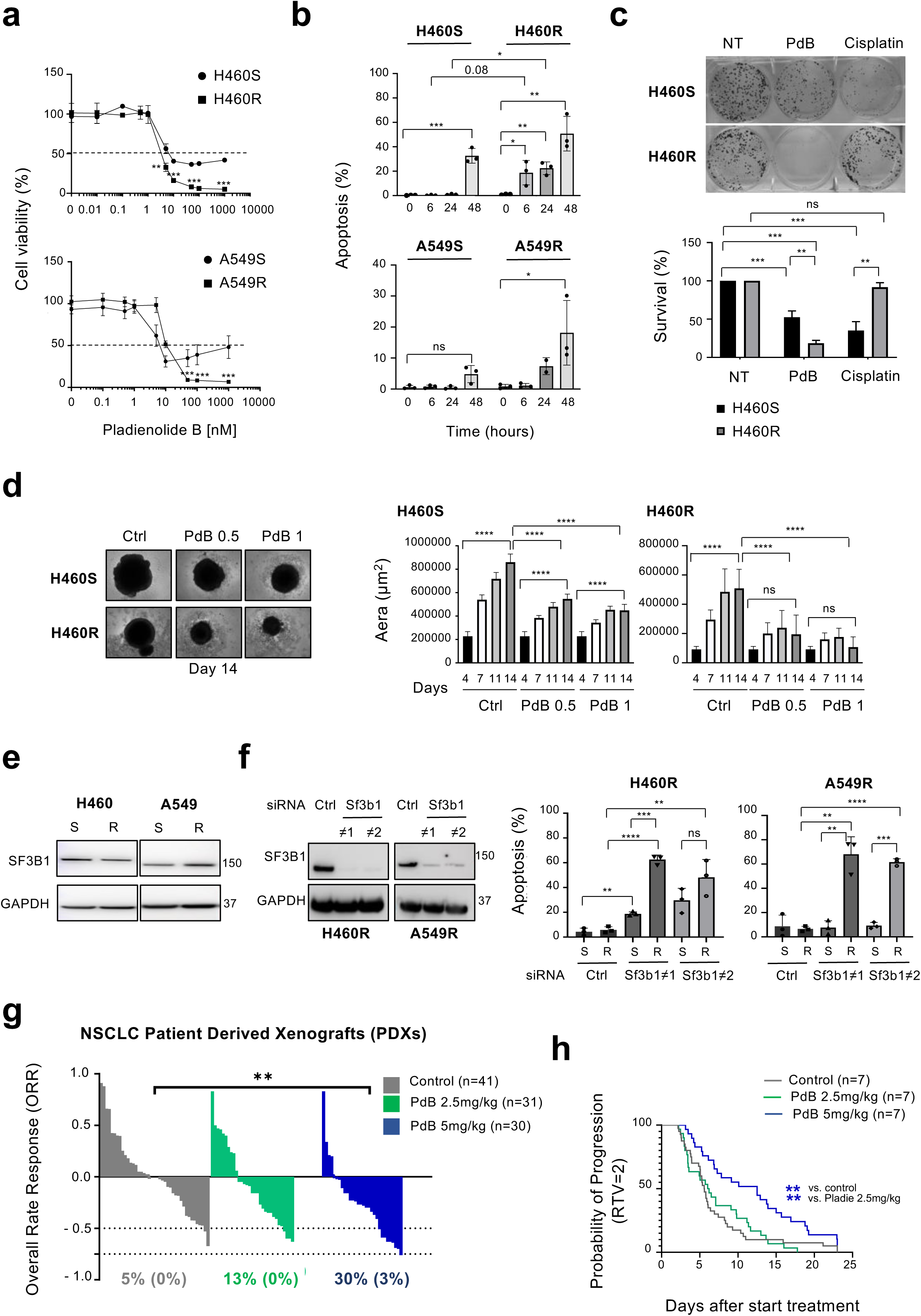
Pladienolide B decreases cell viability in NSCLC cells with acquired resistance to platinum salts and slows down tumor growth in NSCLC Patient Derived Xenografts (PDXs). (a) Cell viability assay in cells treated or not for 72 hours with increasing concentrations of pladienolide B (nM). Mean ± SD. n = 3. (b) Quantification of apoptosis (%) by flow cytometry in cells treated with 5 nM pladienolide B for indicated times. Mean ± SD. n = 3. (c) Clonogenic assay in H460S/R cells treated or not for 14 days with 1 nM pladienolide B or 1 µM cisplatin. Lower panel: number of colonies with number in untreated condition being arbitrarily assigned to 100% survival. Mean ± SD. n = 3. (d) Spheroids from H460S/R cells treated with complete medium and DMSO as control, or 0.5 nM or 1 nM pladienolide B for the indicated days. Upper panels: representative images of spheroids at day 14. Lower panel: spheroids area (µm^2^). Mean ± SD. n = 5 spheroids/condition. A representative experiment of 3 independent ones. (e) Representative immunoblots of SF3B1 in H460S/R or A549S/R cells. GAPDH was used as a loading control. n = 3. (f) H460R or A549R cells were transfected for 72 hours with a control siRNA or two distinct siRNA targeting *Sf3b1* mRNA as indicated. Left panels: representative immunoblots of SF3B1. GAPDH was used as a loading control. Right panels: quantification of apoptosis (%). Mean ± SD. n = 3. (a, b, c, d, f) Unpaired t test. * p< 0.05, ** p< 0.01, *** p< 0.001, **** p< 0.0001, ns: not significant. (g) Overall Rate Response (ORR) at Day 20 for all NSCLC PDXs treated or not (control) with 2.5 mg/kg or 5 mg/kg pladienolide B (ip, Day 1-Day 4-Day 8-Day 11). Two-tails Mann-Whitney t test. ** p< 0.01. (h) Probability of progression (Relative Tumor Volume = 2) in control and pladienolide B-treated PDXs. n = 7 (LCIM1 is not included). Log-rank t test. ** p < 0.01.

**Table 1.**
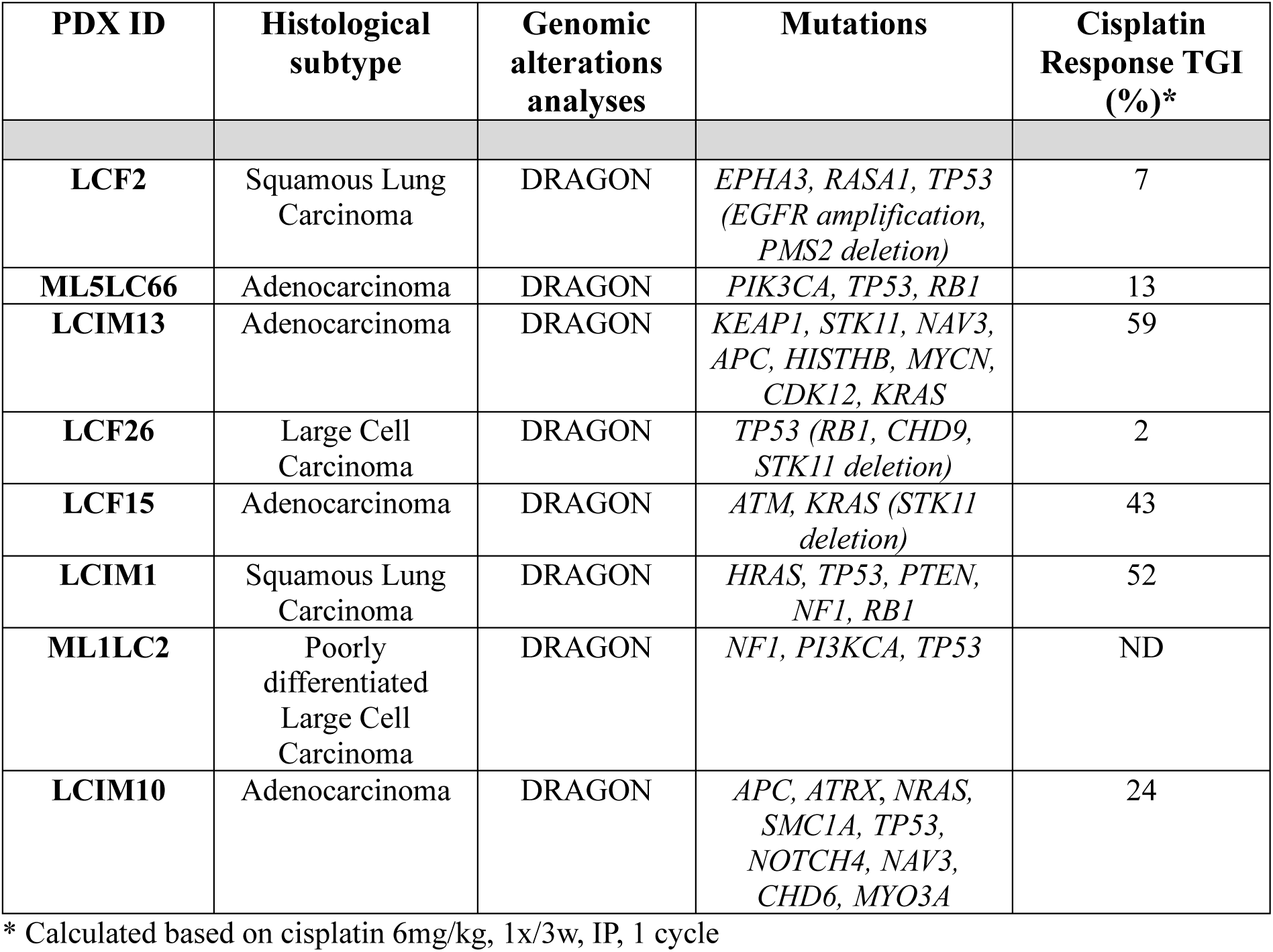
Characteristics (histological subtype, mutations and cisplatin response) of each Non Small Cell Lung Patient-Derived Xenograft used in this study. TGI: Tumor Growth Inhibition. Of note, the LCIM10 PDX which is the most responsive to pladienolide B alone displays ATRX mutation.

### Pladienolide B induces an early replicative stress in NSCLC cells

To deepen the molecular mechanisms involved in the effects of pladienolide B, we first analyze cell cycle distribution. After 24 hours treatment, pladienolide B increased the percentage of H460S/R and A549R cells accumulating in G2/M phase and decreased the percentage of A549S/R cells in S phase (Fig 2a). Pladienolide B also decreased the percentage of H460R cells incorporating EdU, an analog of thymidine, after 6 hours treatment (Fig 2b). These results suggested that pladienolide B may induce replicative stress in NSCLC cells, as previously shown in myelodysplastic syndromes (52). Consistently, a 6 hours treatment increased the number of RPA32 foci/nucleus, a marker of single strand DNAs (ssDNA) (Fig 2c), as well as the number of γH2AX (Fig 2d) and phospho-53BP1(Ser1778) (Fig 2e) foci/nucleus, two markers of DNA double-strand breaks (DSBs). Accumulation of γH2AX upon pladienolide B treatment was confirmed by immunoblotting (Fig 2f). At this time, all these effects were less prominent in H460S cells, consistent with H460R cells exhibiting higher vulnerability to pladienolide B. To go further, we performed *“in situ”* analysis of protein interactions at DNA replication forks (SIRF) that allows to quantify the recruitment of proteins at nascent replication forks (45). We analyzed the recruitment of the DNA polymerase processivity factor PCNA (Proliferating Cell Nuclear Antigen), a marker of active replication forks. We showed that 6 hours pladienolide B treatment decreases the amount of PCNA recruited at nascent replicative forks in H460R cells (Fig 2g). This is consistent with enhanced replicative stress and a slow-down of replication forks (53). Of note, pladienolide B did not decrease total PCNA protein level (Fig 2h). Altogether, these results demonstrated that pladienolide B induces an early replicative stress, notably in NSCLC platinum-salts resistant cells.

**Figure 2.**
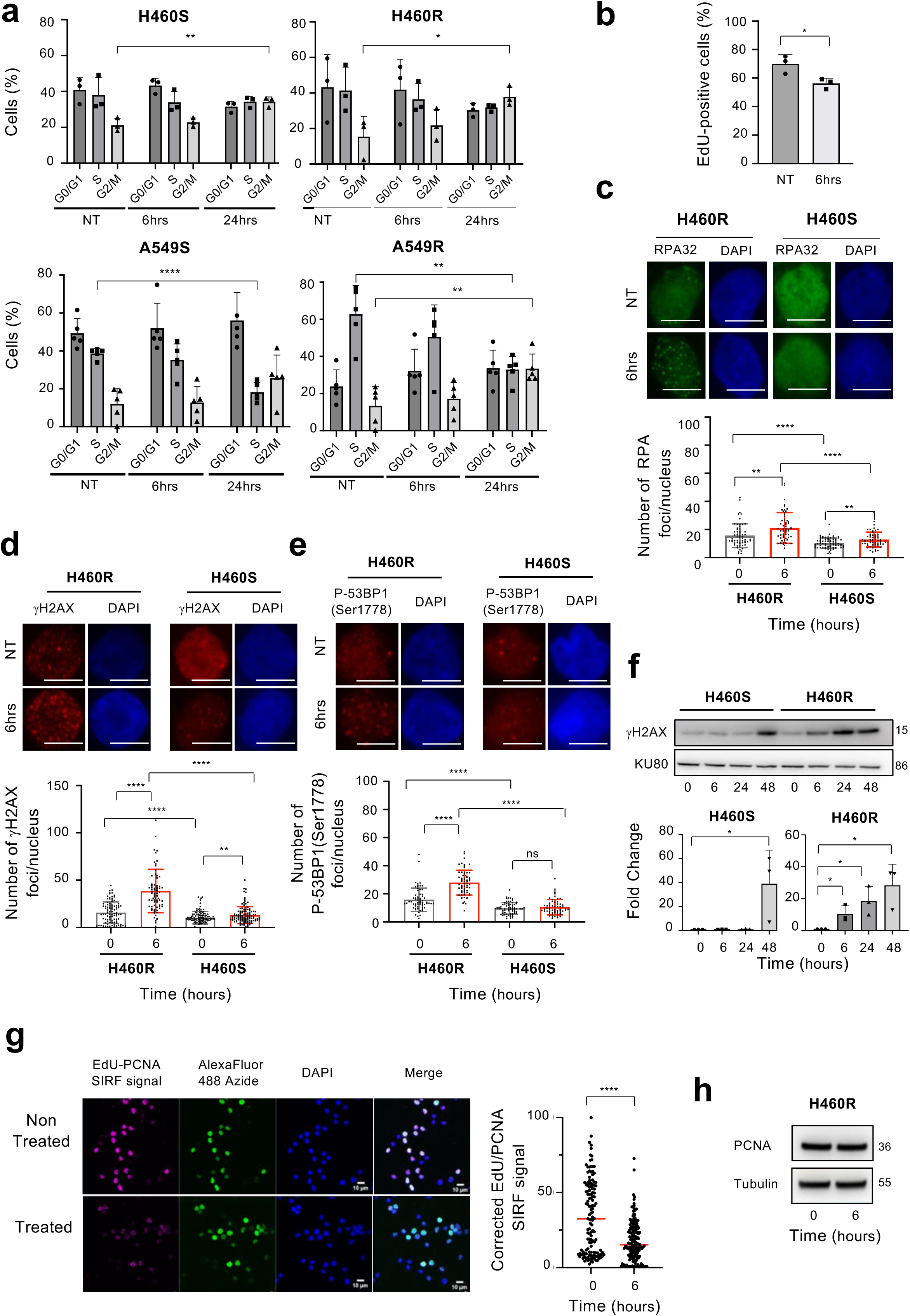
Pladienolide B induces an early replicative stress. (a) H460S/R or A549S/R cells were treated or not with 5 nM pladienolide B for indicated times. Cell cycle distribution (%) in the different phases of the cell cycle. Mean ± SD. n = 3. Unpaired t test. * p< 0.05, ** p< 0.01, **** p< 0.0001. (b) Percentage of EdU-positive H460R cells in untreated or pladienolide B condition. Mean ± SD. n = 3. Unpaired t test. * p< 0.05. (c, d, e) H460R and H460S cells were treated or not with 5 nM pladienolide B for 6 hours. Upper panels: representative immunofluorescence stainings of RPA32 (c), γH2AX (d), or phospho-53BP1(Ser1778) (e). DAPI was used to counterstain the nucleus. Scale bar = 10 µm. Lower panels: quantification of the number of RPA32 (c), γH2AX (d), or phospho-53BP1(Ser1778) (e) foci/nucleus. Mean ± SD. n = 3. Unpaired t test. ** p< 0.01, *** p< 0.001, **** p<0.0001. (f) Upper panels: representative immunoblots of γH2AX protein in H460S/R cells treated with 5 nM pladienolide B for indicated times. KU80 was used as a loading control. Lower panels: densitometric quantification (fold change) of γH2AX signal normalized to KU80 signal. Mean ± SD. n=3. Mann-Whitney t test. * p< 0.05. (g, h) H460R cells were treated or not for 6 hours with 5 nM pladienolide B. (g) SIRF analysis of PCNA recruitment at nascent replication forks. Lefts panels: representative images. Right panel: quantification of EdU-PCNA fluorescent intensity (ratio signal intensity/area)/nucleus. Mean. n = 3. Unpaired t test. **** p<0.0001. (h) PCNA immunoblot. Tubulin was used as a loading control.

### Pladienolide B regulates ATR/DNA-PKcs signaling pathways in NSCLC cells in a transcription-dependent manner

ATR and DNA-PKcs are the main kinases activated in response to replicative stress (54, 55). In H460R and A549R cells mainly, we observed that a 6 hours treatment leads to activation of DNA-PKcs, as reflected by the accumulation of its auto-phosphorylated form P-DNA-PKcs(Ser2056), together with its target P-RPA32(Ser4/8) (Fig 3a-b). In contrast, the active auto-phosphorylated form of ATR, namely P-ATR(Thr1989), did not accumulate upon pladienolide B treatment whatever the cell lines (Fig 3a-b). Replicative stress can be the result of conflicts between replication and transcription machineries (56–58). Therefore, to assess the role of transcription in pladienolide B effects, we used DRB, a selective inhibitor of transcription elongation by RNA polymerase II. As observed by immunoblotting (Fig 3c) or immunofluorescence (Fig 3d), DRB reversed P-DNA-PKcs(Ser2056), P-RPA32(Ser4/8) and/or γH2AX accumulation upon pladienolide B treatment. This suggested that active transcription could account for pladienolide B-induced replicative stress. Then, we assessed the effects of pladienolide B on DNA-PKcs/ATR signaling pathways at later time points (i.e. after 24 and 48 hours exposure). Immunoblotting (Fig 3e) and RT-qPCR (Fig 3f) experiments demonstrated that ATR and DNA-PKcs protein and/or mRNA levels decreased in all cellular models. As a whole, we demonstrated that pladienolide B induces an early transcription-dependent replicative stress that correlates with activation of DNA-PKcs, followed by a shutdown of ATR and DNA-PKcs signaling.

**Figure 3.**
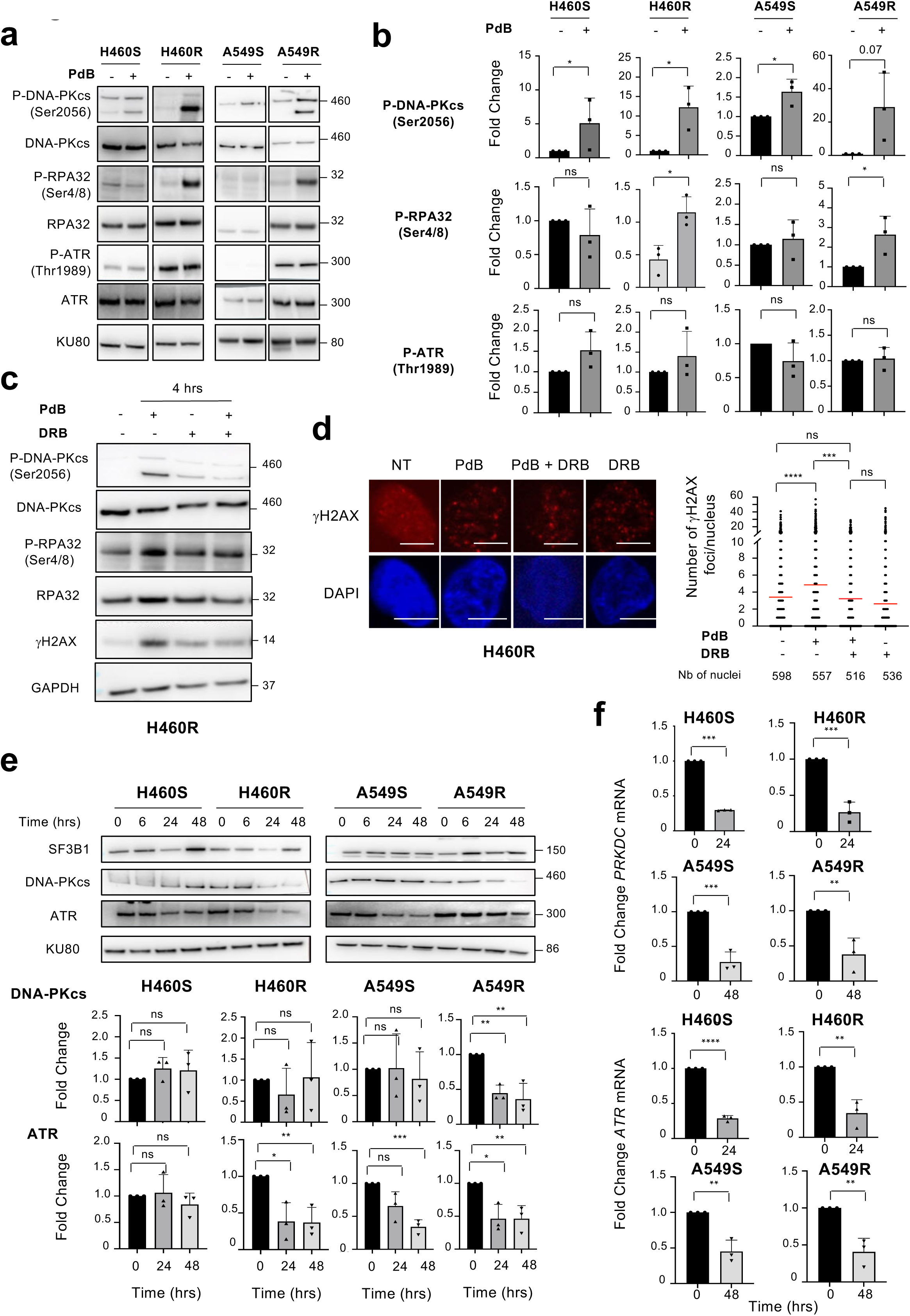
Pladienolide B regulates ATR/DNA-PKcs signaling pathways in a transcription-dependent manner. (a) Immunoblots of the indicated proteins in H460S/R or A549S/R cells treated or not with 5 nM pladienolide B for 6 hours. KU80 was used as a loading control. (b) Densitometric quantification (fold change) of P-DNA-PKcs(Ser2056), P-RPA32(Ser4/8) or P-ATR(Thr1989) signal normalized to KU80 signal. Mean ± SD. n=3. (c) H460R cells were co-treated (+) or not (-) with 5 nM pladienolide B for 4 hours in the presence or absence of 25 µM DRB. Representative immunoblots of the indicated proteins. GAPDH was used as a loading control. n = 3. (d) H460R cells were co-treated (+) or not (-) with 5 nM pladienolide B for 6 hours in the presence or absence of 25 µM DRB. Left panels: representative immunofluorescence stainings of γH2AX in each condition. DAPI was used to counterstain the nucleus. Scale bar = 10 µm. Right panel: quantification of the number of γH2AX foci/nucleus. Mean. n = 3. (e, f) H460S/R or A549S/R cells were treated or not with 5 nM pladienolide B for the indicated times. (e) Upper panels: Immunoblots of the indicated proteins. KU80 was used as a loading control. Lower panels: densitometric quantification (fold change) of DNA-PKcs or ATR signal normalized to KU80 signal. Mean ± SD. n=3. (f) RT-qPCR (fold change) of *PRKDC* or *ATR* mRNA level. Mean ± SD. n = 3. All, unpaired t test. * p< 0.05, ** p< 0.01, *** p< 0.001, **** p<0.0001, ns: not significant.

### SF3B1 knock-down recapitulates the effects of pladienolide B on ATR/DNA-PKcs signaling pathways

To verify that the effects of pladienolide B were not “off-target” effects, we silenced SF3B1 expression using two distinct siRNAs (Fig 4a). We showed that SF3B1 knock-down significantly decreases DNA-PKcs and ATR protein levels in both H460R and A549R cells (Fig 4a). In H460R cells, this decrease was associated with a decrease of *ATR* and *PRKDC* mRNA levels, while this was less the case in A549R cells (Fig 4b). As a whole, these results indicated that SF3B1 silencing recapitulates the effects of pladienolide B on ATR and DNA-PKcs. These results also suggested a close link between SF3B1 and ATR and/or DNA-PKcs in NSCLC. Consistently, looking for correlation between SF3B1 and ATR or DNA-PKcs protein levels among 77 lung cancer cell lines, using quantitative proteomics of the Cancer Cell Line Encyclopedia database (59), we found a significant positive correlation between SF3B1 and DNA-PKcs (r=0.6045, p<0.0001) or ATR (r=0.2336, p=0.0437) protein levels (Fig 4c). Such a positive correlation was also found between *SF3B1* and *PRKDC* (encoding DNA-PKcs) mRNA levels in lung adenocarcinoma patients retrieved from TCGA database (Fig 4d). As a whole, these results indicated that pladienolide B or SF3B1 knock-down negatively impacts DNA-PKcs/ATR signaling pathways, which correlates with cell growth inhibition in NSCLC cell lines.

**Figure 4.**
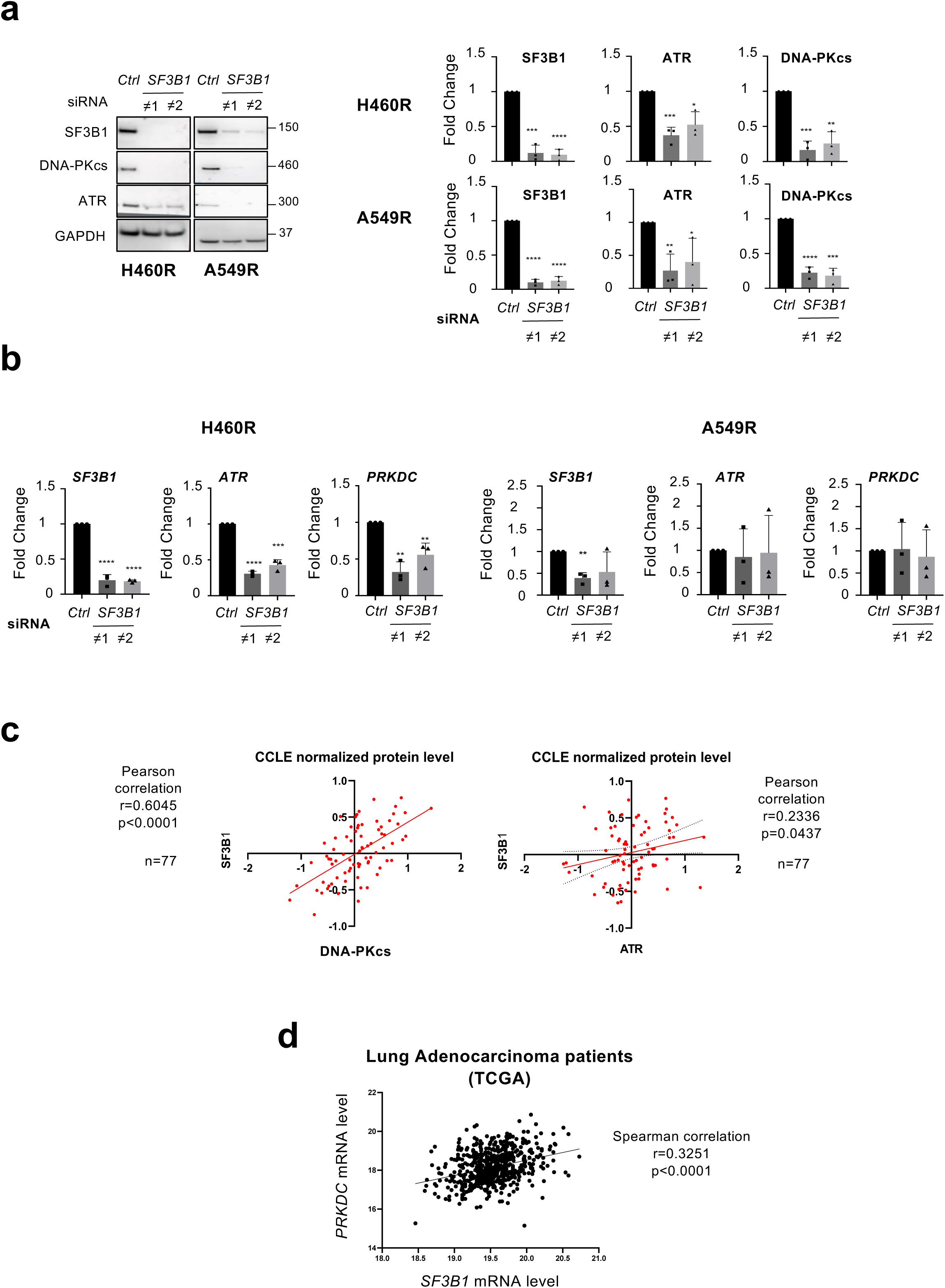
SF3B1 silencing recapitulates the effects of pladienolide B on ATR/DNA-PKcs signaling. (a, b) H460R or A549R cells were transfected for 72 hours with a control siRNA or two distinct siRNA targeting *Sf3b1* mRNA as indicated. (a) Left panels: representative immunoblots of the indicated proteins. GAPDH was used as a loading control. Right panels: densitometric quantification (fold change) of SF3B1, ATR or DNA-PKcs signal normalized to GAPDH signal. Mean ± SD. n=3. (b) RT-qPCR (fold change) of *SF3B1*, *ATR* or *PRKDC* mRNA level. Mean ± SD. n = 3. (a, b) Mann-Whitney t test. * p< 0.05, ** p< 0.01, *** p< 0.001, **** p<0.0001. (c) Correlation between SF3B1 and DNA-PKcs or ATR protein levels in a series of 77 NSCLC cell lines based on CCLE database. (d) Correlation between *SF3B1* and *PRKDC* mRNA levels in lung adenocarcinoma patients retrieved from TCGA public database.

### Pladienolide B massively impacts the expression and splicing of genes involved in DNA damage signaling and repair

At a larger scale, we also investigated the impact of pladienolide B on transcription and splicing in H460S and H460R cells by performing bulk RNA-Seq. Cells were treated with 3 nM pladienolide B for 8 hours before total RNA extraction. Considering Differentially Expressed Genes (DEGs, absolute log2FC ≥ 0.4, p ≤ 0.05), pladienolide B differentially regulated 5453 and 5486 genes in H460S and H460R cells respectively, with 3980 genes being commonly regulated in both cellular models (Fig 5a left panel, Table 2). The number of down-regulated genes was slightly higher compared to that of up-regulated genes (Fig 5a, right panel). This was more pronounced when only DEGs with absolute log2FC ≥ 1 were considered. Hence, pladienolide B decreased the expression of 1036 and 923 genes while increased that of 447 and 471 genes in H460S and H460R cells, respectively (Fig 5b). Interestingly, among these down-regulated genes, Gene Ontology analysis using metascape (48) showed an enrichment in genes involved in DNA metabolic process (GO term:0006259) that includes DNA repair, replication and recombination processes (Fig S3). Thus, to go further, we restricted our analysis to genes belonging to the DNA repair pathways full network (WikiPathways WP4946) including 121 genes as a total. Based on our RNA-Seq data, pladienolide B was found to down-regulate the expression of 41 (33%) and 47 (39%) genes of this network in H460S and H460R cells, respectively (Table 3, absolute log2FC ≥ 0.4, p ≤ 0.05). When DEGs with absolute log2FC ≥ 1 were considered only, we observed decreased expression of genes involved in Mismatch Repair (MMR) (e.g. *POLD3*, *POLE2*, *RFC1*, *RFC4*), Nucleotide Excision Repair (NER) (e.g. *POLD3*, *POLE2*, *GTF2H3*, *RFC1*, *RFC4*), Base Excision Repair (BER) (e.g. *POLD3*, *POLE2*, *PARP2*, *UNG*, *PCNA*), Fanconia Anemia Pathway (FA) (e.g. *FANCE*, *FANCL*), or Homologous Recombination (HR) (e.g. *POLD3*, *RAD51*, *BRCA1*, *BRCA2*) in H460S and/or H460R cells (Fig 5c, Table 3). Hence, pladienolide B downregulates the expression of numerous DNA repair genes in NSCLC cells.

**Figure 5.**
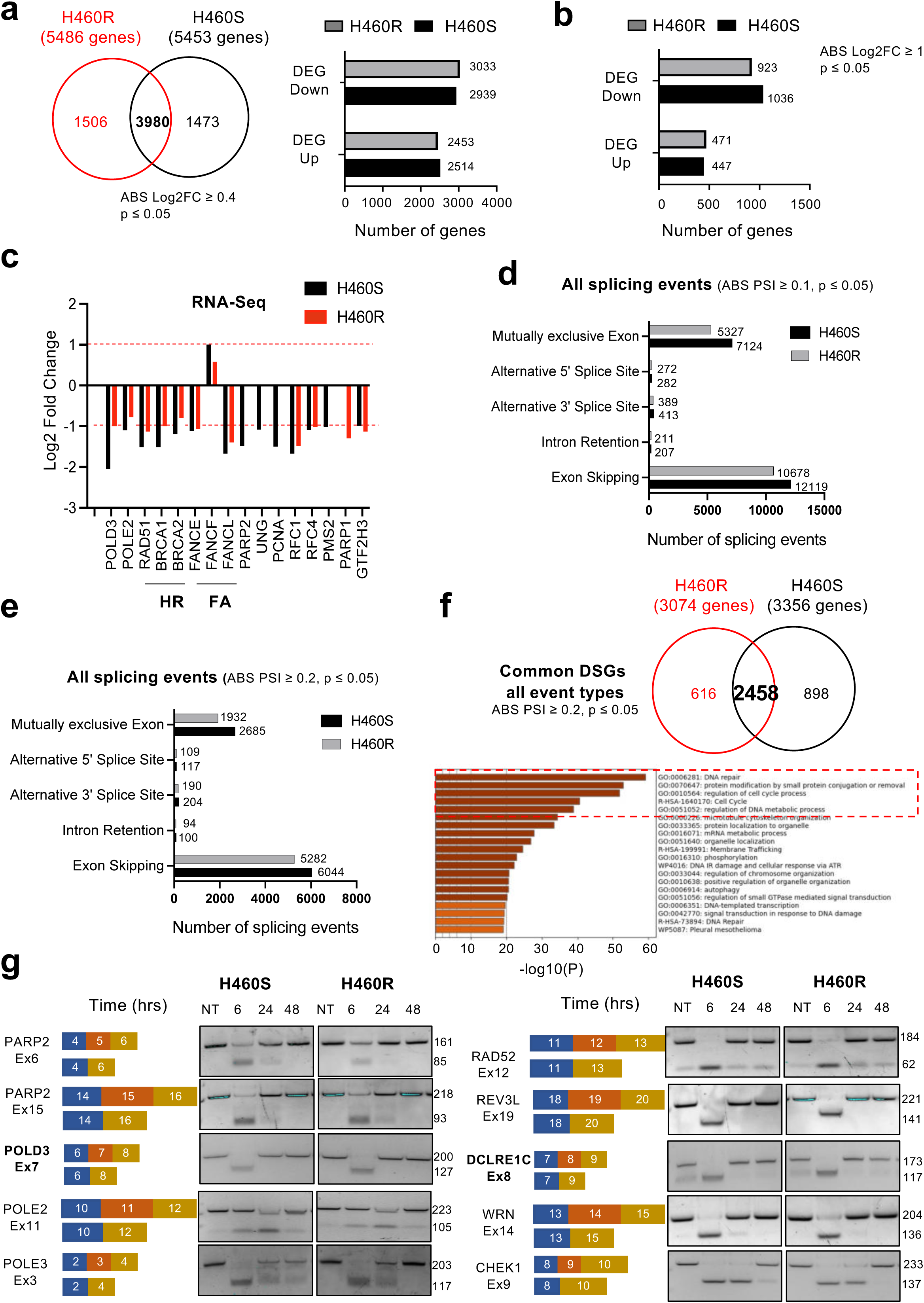
Pladienolide B massively impacts the expression and splicing of genes involved in DNA damage signaling and repair. (a) Left panel: total number of differentially Expressed Genes (DEGs) based on RNA-Seq studies in H460S/R cells treated with 3 nM pladienolide B for 8 hours as compared to untreated condition. Genes with p value ≤ 0.05 and absolute Log2 FC ≥ 0.4 were considered as significant. Right panel: number of DEGs either down- or up-regulated. (b) Number of DEGs either down- or up-regulated in H460S/R cells according to absolute Log2 FC ≥ 1 and p value ≤ 0.05. (c) DEGs belonging to the DNA repair pathways full network (WikiPathways, WP4946) in H460S/R cells. (d, e) Number of total splicing events, according to each type of alternative splicing, differentially regulated in H460 parental or resistant cells treated with 3 nM pladienolide B for 8 hours as compared to untreated condition. (d) Number of differential splicing events with p value ≤ 0.05 and absolute deltaPSI ≥ 10%. (e) Number of differential splicing events with p value ≤ 0.05 and absolute deltaPSI ≥ 20%. (f) Upper panel: number of specific and common Differentially Spliced Genes (DSGs) with absolute deltaPSI ≥ 20% in H460R and H460S cells. Lower panel: Gene Ontology Analysis on common DSGs (n = 2458). (g) RT-PCR validation of exon skipping events impacting genes involved in DNA repair pathways in H460S/R cells treated or not with 5 nM pladienolide B for indicated times. Representative agarose gels of amplified products. Numbers indicate amplicon size (in base pairs). n = 3.

**Table 2.** Differentially Expressed Genes (DEGs) in H460 parental or resistant cells treated with 3 nM pladienolide B for 8 hours as compared to untreated cells. p value ≤ 0.05, absolute log2FC ≥ 0.4.

**Table 3.** Differentially Expressed Genes (DEGs) belonging to the DNA repair pathways full network (WikiPathways, WP4946) in H460 parental or resistant cells treated with 3 nM pladienolide B for 8 hours as compared to untreated cells. p value ≤ 0.05, absolute log2FC ≥ 0.4. Gene name, log2 fold change, down- or up-regulation, and type of DNA repair in which genes are involved are listed. In bold, DEG with p value ≤ 0.05 and absolute log2FC ≥ 1.

We also looked at splicing events. Considering all differential splicing events [Absolute Percent Splice In (PSI) ≥ 0.1, p ≤ 0.05], and according to each type of alternative splicing (exon skipping, intron retention, alternative 3’SS, alternative 5’SS, mutually exclusive exon), pladienolide B was found to mostly regulate exon skipping and mutually exclusive exon in both cellular models (Fig 5d, Table 4). Being more stringent, and considering splicing events with absolute PSI ≥ 0.2 only, we found that pladienolide B regulates 6044 and 5282 exon skipping events, 2685 and 1932 mutually exclusive exons, 117 and 109 alternative 5’SS, 204 and 190 alternative 3’SS as well as 100 and 94 intron retention events in H460S and H460R cells, respectively (Fig 5e, Table 4). All these splicing events corresponded to 3356 and 3074 genes that were differentially spliced in H460S and H460R cells respectively, with 2458 genes being commonly regulated in both cellular models (Fig 5f). When Gene Ontology analysis was done in H460S or H460R cells according to each type of alternative splicing, we observed an enrichment in genes involved in DNA Damage Response or DNA metabolic process among genes with exon skipping or mutually exclusive exon (Fig S4). In contrast, we did not see any enrichment for DNA damage-involved genes among other alternative splicing types such as intron retention, 3’ASS or 5’ASS. Importantly, crossing common Differentially Splice Genes between H460S and R cells led to a higher significant enrichment in genes involved in DNA repair (Fig 5f). Crossing also Differentially Splice Genes (absolute PSI ≥ 0.2) with Differentially Expressed Genes (absolute log2FC ≥ 0.4) in either H460S (Fig S5a) or H460R (Fig S5b) cells led to the identification of a list of 1560 and 1496 genes commonly regulated respectively, and enriched in GO terms related to DNA Damage Response. Therefore, beyond transcriptional regulation, these results highly support the idea that deregulated RNA splicing also accounts to the global effects of pladienolide B on DNA damage signaling and repair. To validate some of these splicing events by RT-PCR, we focused on exon skipping events as they were the most prominent events in both cellular models. Initially, we selected 12 exon skipping events on 9 genes, namely *MLH3*, *MSH5*, *PNKP*, *RAD54L*, *EME1*, *DCLRE1C*, *POLD3*, *SETMAR* and *SMC6*, as they were predicted to occur only in H460R cells upon pladienolide B treatment based on RNA-Seq data. Four of these exon skipping events were not validated (*PNKP*-Ex9, *DCLRE1C*-Ex11, *SETMAR*-Ex2, *SMC6*-Ex6). For *MSH5*-Ex17, *RAD54L*-Ex7, *RAD54L*-Ex15, *EME1*-Ex4, *DCLRE1C*-Ex8 and *POLD3*-Ex7, we validated exon skipping with no clear difference between H460R and H460S cells (Fig 5g and Fig S6). Interestingly, most of these exon skipping events occurred early, following 6 hours pladienolide B treatment, and recovered at baseline level 24 to 48 hours after the beginning of treatment. This could be reminiscent with the recent work of Caggiano et al. who demonstrated in triple negative breast cancer cells that a pulse of splicing inhibition is sufficient to trigger a long lasting effect on gene involved in DNA repair, leading to persistent DNA damage (60). To extend the list of our validations, we again focused on genes belonging to DNA repair pathways full network (WikiPathways, WP4946) and looked at exon skipping events that could be regulated in H460S or H460R cells. We found 107 and 87 exon skipping events affecting 59 and 48 genes of this network involved in all DNA repair pathways in H460S and H460R cells, respectively (Table 5). 45 genes were predicted to be commonly spliced in both H460S and H460R cells. As illustrated examples of RT-PCR, we validated skipping of *APEX2*-Ex3, *PARP2*-Ex6, *PARP2*-Ex15, *POLE*-Ex18, *POLE2*-Ex11, *POLE3*-Ex3, *RAD52*-Ex12, *LIG1*-Ex3, *CUL4A*-Ex21 or *GTF2H3*-Ex4 in both cellular models (Fig 5g and Fig S6). Again, most of these skipping exons occurred at earlier time point and did not persist whatever the cellular model. As a whole, these results identified a massive, but only transient, impact of pladienolide B on exon skipping of numerous genes involved in various DNA repair pathways.

**Table 4.** Differentially Spliced Genes (DSGs), according to each type of alternative splicing, in H460 parental or resistant cells treated with 3 nM pladienolide B for 8 hours as compared to untreated cells. ES: exon skipping; MXE: mutually exclusive exon; IR: intron retention; A3SS: alternative 3’ splice site; A5SS: alternative 5’ splice site. p value ≤ 0.05, absolute delta Percent Splice In (PSI) ≥ 0.1.

**Table 5.** Differential exon skipping events affecting genes belonging to the DNA repair pathways full network (WikiPathways, WP4946) in H460 parental or resistant cells treated with 3 nM pladienolide B for 8 hours as compared to untreated cells. For each gene, delta PSI, exon number and DNA repair pathways in which gene is involved are highlighted. p value ≤ 0.05, absolute delta Percent Splice In (PSI) ≥ 0.1.

### Pladienolide B leads to a global impairment of DNA double-strand breaks (DSB) repair pathways which correlates with increased genomic instability in NSCLC cells

The massive effect of pladienolide B on the expression and/or splicing of numerous genes involved in DNA damage signaling and repair pushed us to investigate whether pladienolide B negatively impacts the DNA repair capacities of NSCLC cells. To answer, we took advantage of two cellular models recently engineered in the laboratory allowing to assess the repair of DNA double-strand breaks (DSBs), either by homologous recombination (HR) or canonical Non-Homologous End-Joining (c-NHEJ) (43). H1299-pBL174 cells are derived from H1299 NSCLC cells and stably express the pBL174-pDR-GFP plasmid. This plasmid allows the monitoring of HR repair of a DSB induced by the nuclease I-SceI, upon transient transfection with the pBL133-encoding-I-SceI plasmid, through the re-creation of a functional GFP (Fig 6a and see materials and methods). In pBL133-tranfected cells, pladienolide B significantly decreased the percentage of GFP-positive cells as compared to untreated cells (Fig 6b). To investigate whether pladienolide B also regulates c-NHEJ, we used stable clones derived from the A549 NSCLC cells and obtained after transfection with the pBL230 plasmid that allows to monitor repair of I-SceI-induced DSBs by c-NHEJ through the expression of CD4 (Fig 6c and see materials and methods) (43). Pladienolide B did not impact the expression level of I-SceI following transient transfection with the pBL133 plasmid (Fig 6d, left panel). However, it decreased the percentage of CD4-positive cells following 24 or 48 hours treatment (Fig 6d, right panel). In agreement with a loss of DNA DSB repair capacities, pladienolide B increased chromosomal breaks and exchanges in H460R cells after 48 hours treatment (Fig 6e). Altogether, these results demonstrated that pladienolide B inhibits DNA DSB repair in NSCLC cell lines which might contribute to enhanced genomic instability. Importantly, in clonogenic survival assays, combining pladienolide B with pharmacological inhibitors targeting crucial components of DSB repair, namely RAD51, RAD52 or PARP1/2, strongly increased cell growth inhibition compared to each treatment alone in both H460S and H460R cells (Fig 6f). These results provide the rationale for combining pladienolide B with drugs inhibiting various DNA repair pathways to increase cell death in NSCLC cells.

**Figure 6.**
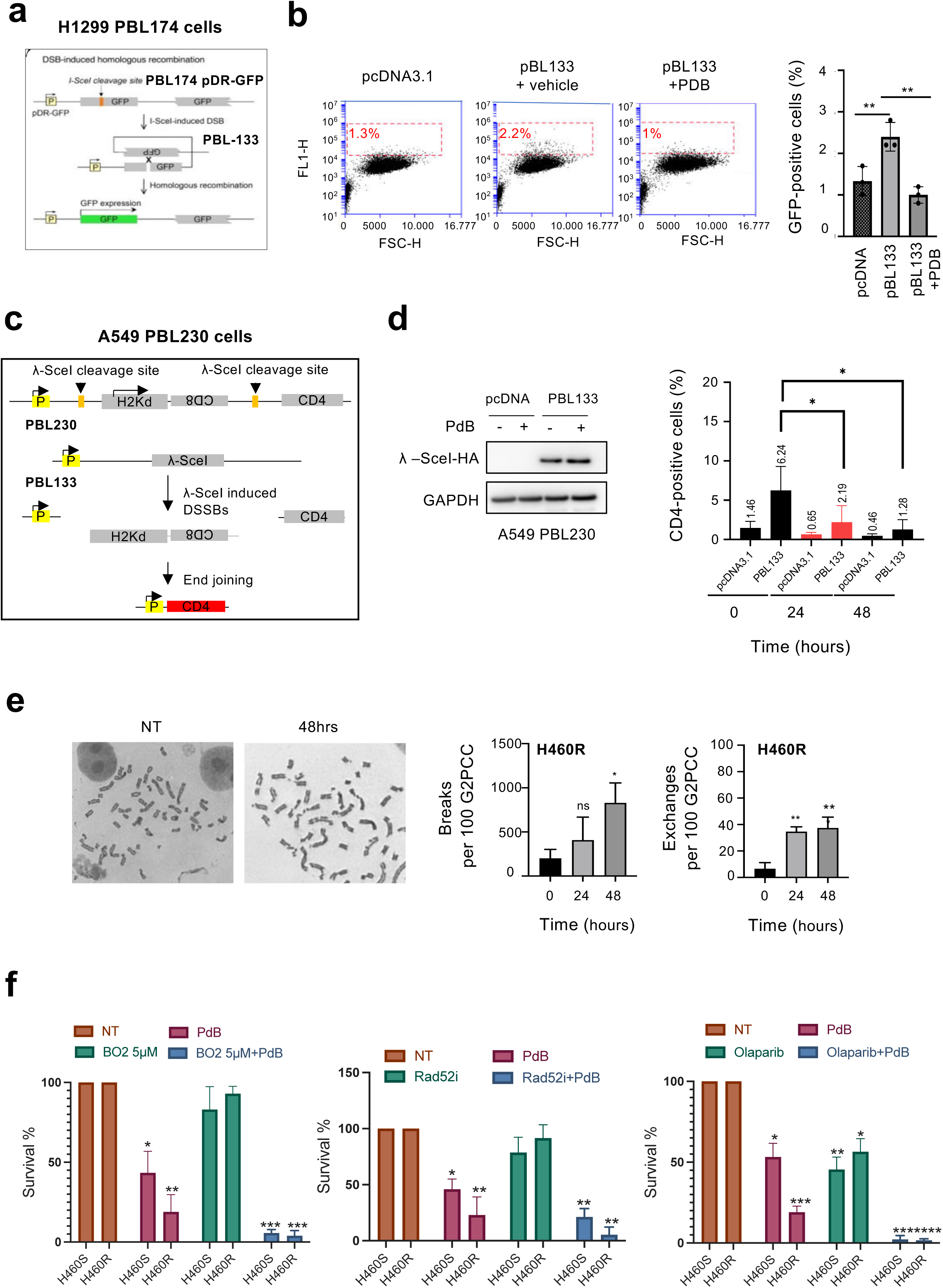
Pladienolide B inhibits DNA double-strand breaks (DSB) repair, induces genomic instability and exhibits enhanced cytotoxicity when combined with drugs targeting distinct DNA DSB repair pathways. (a) Quantification of DSB-induced homologous recombination in H1299 clones stably expressing the pDR-GFP (pBL174) plasmid. When these cells are transfected with the pBL133 plasmid encoding the I-SceI restriction enzyme, efficient recombination restores a functional GFP coding sequence. (b) H1299-pBL174 cells were transfected with pcDNA3.1 or pBL133 for 24 hours and treated (pBL133+PDB) or not (pBL133) with 5 nM pladienolide B during 48 additional hours. Left panels: representative FL1-H versus FSC-H dot plots of GFP-positive (FL1-H +) and –negative (FL1-H -) cells. Right panel: GFP-positive cells (%). Mean ± SD. n = 3. Unpaired t test. ** p<0.01, ns: not significant. (c) A549 LUAD cells stably expressing the pBL230 plasmid were generated. This plasmid contains genes encoding the membrane antigens CD4 and CD8. CD8 is not expressed as it is in inverted orientation, and CD4 is not expressed because it is too far from the promoter. Two cleavage sites for I-SceI are present in non-coding sequences, which are in direct orientation generating cohesive ends between the two sites. When two DSBs are produced by I-SceI, rejoining of the DNA ends by exclusion or inversion leads to the expression of CD4 or CD8, respectively. (d) Quantification of DNA repair by c-NHEJ in A549-pBL230 clones transfected with either pcDNA3.1 or pBL133 plasmid and treated or not with 5 nM pladienolide B for 24 or 48 hours. Left panel: western blotting of I-SceI-HA in transfected cells treated or not with pladienolide B for 48 hours. Right panel: CD4-positive cells (%). Mean ± SD. n = 3. Mann-Whitney t test. * p< 0.05. (e) G2PCC in H460R cells treated or not with 5 nM pladienolide B for indicated times. Left panels: representative images of G2PCC showing chromosome breaks and exchanges. Right panels: quantification of the number of breaks or exchanges per 100 G2PCC. Mann-Whitney t test. * p< 0.05, ** p<0.01, ns: not significant. (f) Clonogenic survival assay. H460S or H460R cells were treated or not with 1 nM pladienolide B for 10 days in the presence or absence of 5 µM BO2 a RAD51 inhibitor, which was the only drug removed after 24 hours treatment, 10 µM 6-OH-DOPA, a RAD52 inhibitor, or 10 µM olaparib, a PARP1/2 inhibitor. The number of colonies obtained in non treated condition was arbitrarily assigned to 100% survival. Mann-Whitney t test. * p< 0.05, ** p<0.01, *** p< 0.001, **** p<0.0001.

### Pladienolide B or SF3B1 controls *MLH3* splicing and protein levels in NSCLC cells

During the RT-PCR validation step of pladienolide B-induced exon skipping events, we identified skipping of *MLH3*-Ex8 as the sole event displaying a slight difference between H460S and H460R cells. Of note, exon 8 was annotated based on FASTERDB. It corresponds to exon 7 based on ENSEMBL (ENSE00003584609). Hence, exon skipping of *MLH3*-Ex8 persisted after 24 hours pladienolide B treatment in both cell lines and full recovery occurred only after 72 hours treatment in H460R cells (Fig 7a, upper panels). The same tendency was observed in A549R cells (Fig 7a, lower panels). Interestingly, *MLH3*-Ex8 encodes the endonuclease domain of MLH3 thereby suggesting that exon skipping could alter MLH3 function (61, 62). Based on RNA-Seq data, *MLH3*-Ex5 was also predicted to be skipped specifically in H460R cells. However, we did not observe significative skipping of *MLH3*-Ex5 upon pladienolide B treatment in the H460 cellular model (Fig 7b, upper panels), while *MLH3*-Ex5 skipping occurred in both A549S and A549R cells and tended to persist until 72 hours in A549R cells (Fig 7b, lower panels). SF3B1 knock-down (Fig 7c, left panels) led to *MLH3*-Ex8 skipping in both H460R and A549R cells while it did not impact *MLH3*-Ex5 skipping (Fig 7c, right panels). Interestingly, SF3B1 neutralization also strongly decreased MLH3 protein level in both H460R and A549R cells (Fig 7d). Altogether, these results identified *MLH3* splicing, and more precisely *MLH3*-Ex8 and/or -Ex5 skipping, as being regulated by both pladienolide B and SF3B1 silencing in NSCLC cells. Interestingly, and consistent with a closed relationship between SF3B1 and MLH3 in NSCLC, we observed a positive correlation between *SF3B1* mRNA level and *MLH3*-Ex8 (r=0.2391,p<0.0001) or MLH3-Ex5 (r=0.2498, p<0001) usage in lung adenocarcinoma patients (Fig 7e). These data are consistent with MLH3 being a bona fide target of SF3B1 in lung cancer.

**Figure 7.**
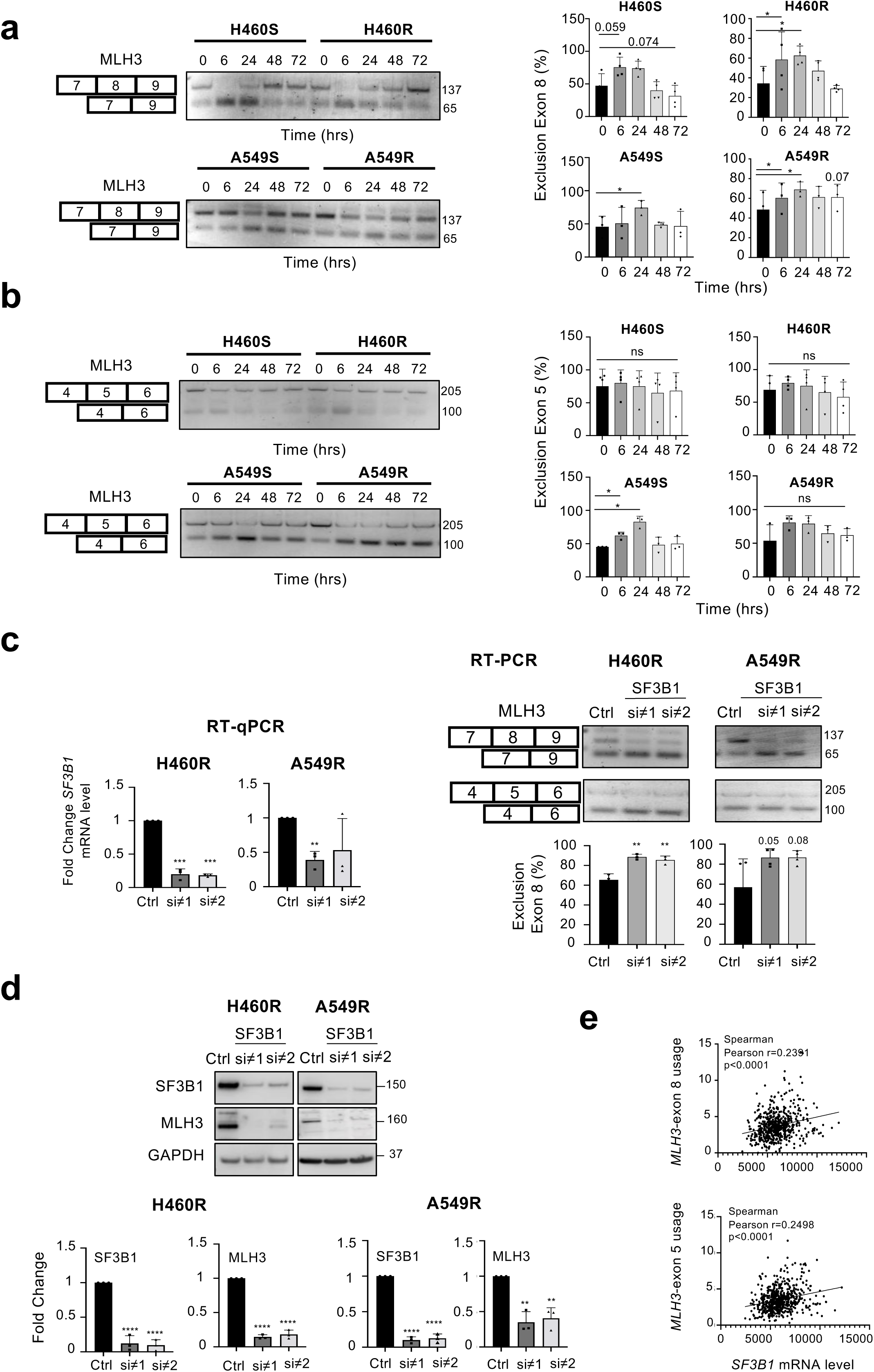
Pladienolide B and/or SF3B1 knock-down induces *MLH3*-Exon 8 skipping and decreases MLH3 protein level. (a, b) H460S/R or A549S/R cells were treated with 5 nM pladienolide B for indicated times. RT-PCR analysis of skipping of *MLH3*-Ex8 (a) or *MLH3*-Ex5 (b). Left panels: representative agarose gels of amplified products. Numbers indicate amplicon size (in base pairs). Right panels: densitometric quantification of exon 8 or 5 exclusion (%). Mean ± SD. n = 4 (H460S/R) or n = 3 (A549S/R). (c, d) H460R or A549R cells were transfected with control siRNA or two distinct siRNAs targeting *SF3B1* for 72 hours. (c) Left panels: RT-qPCR (fold change) of *SF3B1* mRNA level. Mean ± SD. n = 3. Right panels: RT-PCR analysis of *MLH3*-Ex8 or *MLH3*-Ex5 skipping. Upper panels: representative agarose gels of amplified products. Numbers indicate amplicon size (in base pairs). Lower panels: densitometric quantification of exon 8 exclusion (%). Mean ± SD. n = 4. (d) Upper panel: representative immunoblots of SF3B1 or MLH3 protein levels. GAPDH was used as a loading control. Lower panels: densitometric quantification of SF3B1 or MLH3 protein level according to GAPDH. Number obtained in non treated condition was arbitrarily assigned the value 1. Mean ± SD. n = 3. All, Mann-Whitney t test. **p< 0.01, *** p<0.001, **** p<0.0001. (e) Correlation between *SF3B1* mRNA level and usage of *MLH3*-Ex8 or *MLH3*-Ex5 in lung adenocarcinoma patients.

### Pladienolide B counteracts resistance to cisplatin in NSCLC cells which correlates with enhanced skipping of *MLH3*-Ex 8 and decreased DNA-PKcs and MLH3 protein levels

Finally, we wondered whether pladienolide B, by negatively regulating DNA damage response and repair, reverses resistance to platinum salts. To answer, H460R or A549R cells were co-treated with pladienolide B and/or cisplatin for 24 hours and apoptosis was quantified by active-caspase-3 staining followed by flow cytometry. Combining pladienolide B and cisplatin significantly increased cellular apoptosis in both H460R and A549R models (Fig 8a). As detected by immunoblotting, such increase in apoptosis was associated with enhanced accumulation of active caspase-3 and γH2AX, as well as enhanced decrease of DNA-PKcs protein levels (Fig 8b-c). To go further, we analyzed various exon skipping events found to be impacted by pladienolide B alone, including genes involved in Nucleotide Excision Repair (NER) as it is a major pathway involved in the repair of cisplatin-induced DNA adducts (63, 64). These genes included *GTF2H3*-Ex4, *GTF2H4*-Ex9, *Cul4A*-Ex21, *POLE2*-Ex11, *LIG1*-Ex3, *POLD3*-Ex7, or *POLE3*-Ex3. We did not see any differences whatever the condition and time (data not shown). Then, we looked at *MLH3* splicing. As compared to untreated condition and each drug used alone, combining pladienolide B and cisplatin tended to increase *MLH3*-Ex8 skipping in H460R cells, whereas *MLH3*-Ex5 skipping was not affected (Fig 8d-e). In A549R cells, no difference was observed between pladienolide B and pladienolide B combined to cisplatin. *MLH3*-Ex8 was strongly skipped in both conditions compared to untreated cells or cisplatin used alone (Fig 8d-e). As quantified by RT-qPCR, *MLH3* mRNA level also tended to increase in H460R cells treated with pladienolide B or the combination of pladienolide B and cisplatin, while no difference was observed in A549R cells (Fig 8f). Moreover, as SF3B1 silencing, pladienolide B decreased MLH3 protein level both in H460R and A549R cells (Fig 8g), with the combination pladienolide B and cisplatin exhibiting a stronger effect in H460R cells. As a whole, these results demonstrated that pladienolide B counteracts resistance of NSCLC cells with acquired resistance to platinum salts, which correlates with *MLH3*-Ex8 skipping and decrease of DNA-PKcs and MLH3 protein levels.

**Figure 8.**
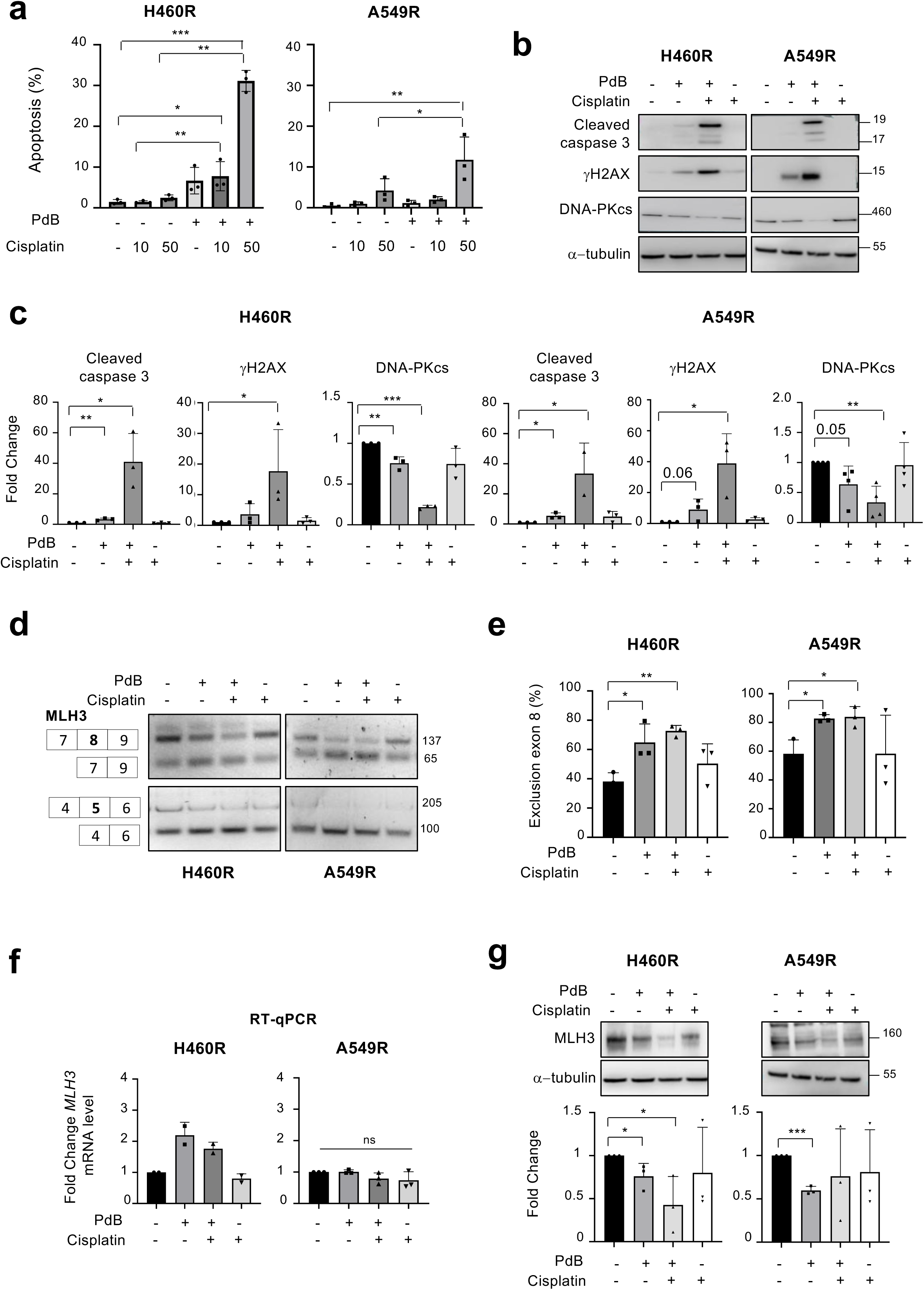
Pladienolide B counteracts resistance to cisplatin in NSCLC cells which correlates with *MLH3*-Ex8 skipping and enhanced decrease of DNA-PKcs and MLH3 protein levels. (a-g) H460R or A549R cells were treated (+) or not (-) for 24 hours with 5 nM pladienolide B in the presence or absence of 10 µM (a) or 50 µM (a, b, c, d, e, f, g) cisplatin as indicated. (a) Quantification of apoptosis (%) by flow cytometry. Mean ± SD. n = 3. (b) Representative immunoblots of indicated proteins. α-tubulin was used as a loading control. n = 3. (c) Densitometric quantification of the signal obtained for the indicated proteins according to the signal obtained with α-tubulin. Number obtained in non treated condition was arbitrarily assigned the value 1. Mean ± SD. n = 3. (d) RT-PCR analysis of skipping of *MLH3*-Ex8 or *MLH3*-Ex5. Representative agarose gels of amplified products. Right numbers indicate amplicon size (in base pairs). (e) Densitometric quantification of exon 8 exclusion (%). Mean ± SD. n = 3. (f) RT-qPCR (fold change) of *MLH3* mRNA level. (g) Upper panels: representative immunoblots of MLH3 protein levels. α-tubulin was used as a loading control. Lower panels: normalized densitometric quantification (fold change) of MLH3 signal according to α-tubulin signal. Number obtained in untreated condition was arbitrarily assigned the value 1. All, Mann-Whitney t test. * p< 0.05, ** p< 0.01, *** p< 0.001, ns: not significant.

### Pladienolide B counteracts resistance to cisplatin in NSCLC PDXs which also correlates with decreased levels of ATR/DNA-PKCs and/or enhanced skipping of *MLH3*-Ex 8

Finally, we wondered whether combining pladienolide B with cisplatin could provide additional therapeutic benefit *“in vivo”*. To answer, we treated NSCLC PDXs either with cisplatin or pladienolide B alone, or a combination of both. We excluded the LCIM10 PDX that was found to be the more responsive to pladienolide B alone (Fig S2b-c). According to the Overall Rate Response (ORR), we found that the combination of pladienolide B and cisplatin enhances tumor growth inhibition compared to each drug alone (Fig 9a). Upon combination, 28% of PDXs reached an ORR< -0.7 compared to 4% of PDXs in each drug alone condition. When we analyzed the probability of progression by calculating RTV=2 (Fig 9b) or RTV=4 (Fig S7a), we observed an enhancement of time to progression upon pladienolide B + cisplatin treatment compared to control or cisplatin alone. When each PDX and either ORR (Fig S7b) or mean RTV (Fig S7c) were considered, we observed that pladienolide B alone or in combination with cisplatin tends to slow down tumor growth in four PDXs, namely LCF26 (pladienolide B alone), ML1LC2, LCIM1 and ML5LC66 (pladienolide B and cisplatin). Immunoblotting experiments provided evidence that pladienolide B, but not cisplatin, decreases DNA-PKcs protein level in two of them (LCF26, ML1LC2) as well as ATR protein level in LCF26 (Fig 9c). In addition, in 3 out of 4 PDXs (LCF26, ML1LC2, LCIM1), RT-PCR analyses demonstrated a slight increase in *MLH3*-exon 8 skipping in response to the combination treatment as compared to each drug alone (Fig 9d). Of note, in these PDXs, we also assessed skipping of *MLH3*-Ex5, *MSH5*-Ex17, *EME*-Ex4, *CHEK1*-Ex9, *EXO1*-Ex5, *POLD3*-Ex7 or *RAD54L*-Ex15, without any variations whatever the treatment condition (data not shown). As a whole, these results provide the first evidence that pladienolide B counteracts resistance to cisplatin in NSCLC PDXs. In addition, they identify ATR, DNA-PKcs and MLH3 as “in vivo” targets of pladienolide B, used alone or in combination with platinum salts. Of note, immunoblotting experiments comparing basal levels of endogenous activated P-ATR(Thr1989) protein among distinct NSCLC PDXs, including the 8 PDXs we studied, showed detectable level in LCF26, LCIM1, ML5LC66, ML1LC2, LCIM10 responsive PDXs while this was not seen for LCF2, LCIM13 or LCF15 non-responsive PDXs (Fig S7d).

**Figure 9.**
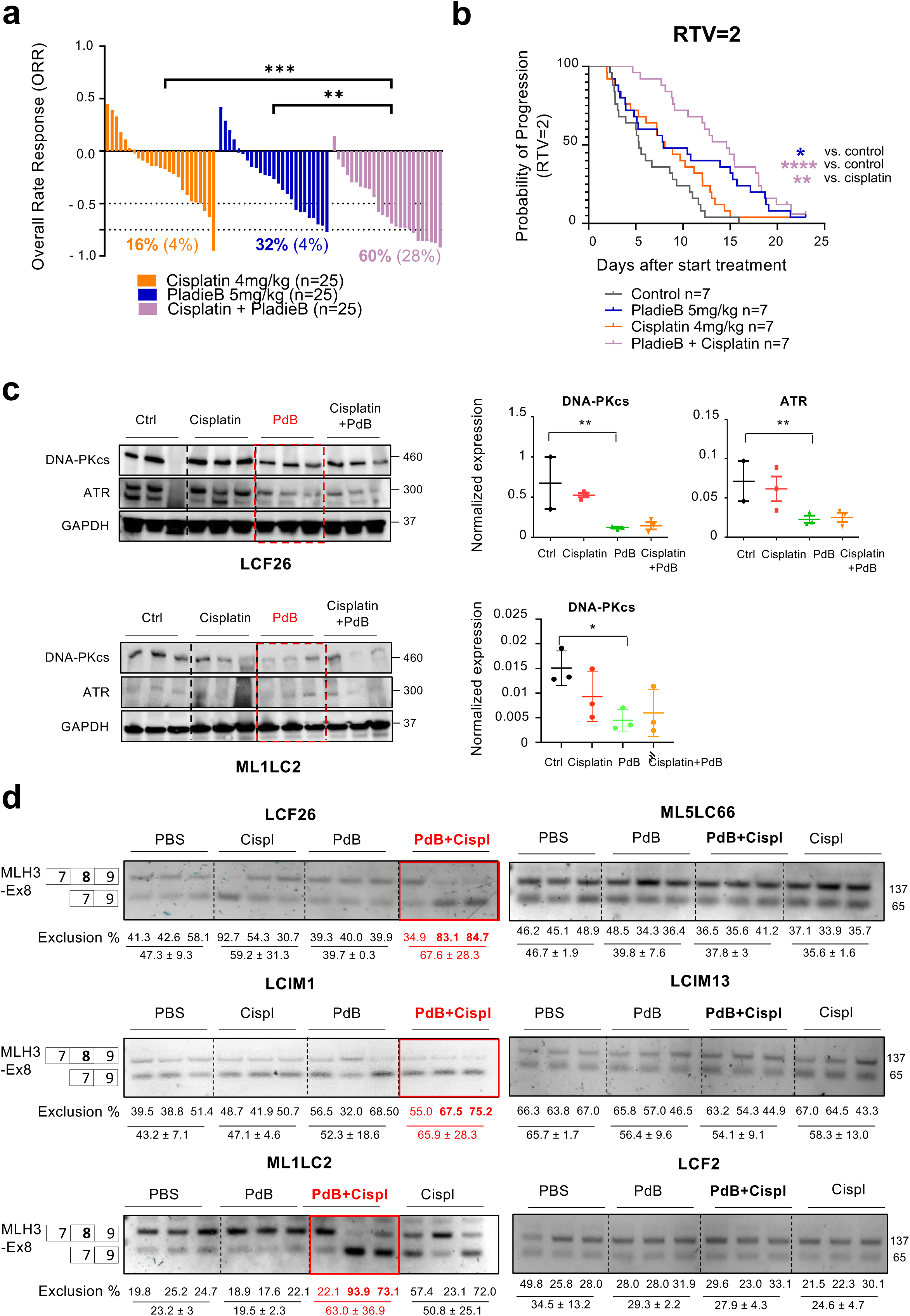
Pladienolide B counteracts resistance to cisplatin in NSCLC PDXs which also correlates with decreased levels of ATR/DNA-PKCs and/or enhanced skipping of *MLH3*-Ex 8. (a) Overall Rate Response (ORR) of all NSCLC PDXs (n = 25) treated with 4 mg/kg cisplatin, 5 mg/kg pladienolide B or a combination of both as compared to vehicle condition. Two-tails Mann-Whitney t test. ** p< 0.01, *** p< 0.001. (b) Probability of tumor progression (Relative Tumor Volume = 2) in control, pladienolide B-, cisplatin- or pladienolide B + cisplatin-treated PDXs. n = 7. Log-rank test. * p< 0.05, ** p < 0.01, *** p< 0.001. (c) LCF26 and ML1LC2 PDXs were treated with vehicle (ctrl), 4 mg/kg cisplatin, 5 mg/kg pladienolide B, or a combination of both. Left panels: immunoblots of DNA-PKCs and ATR. 3 PDXs/condition. GAPDH was used as a loading control. Right panels: normalized expression of DNA-PKcs or ATR according to GAPDH. Mann-Whitney t test. * p< 0.05, ** p< 0.01. (d) RT-PCR analysis of skipping of *MLH3*-Ex8 in each PDX (3 distinct mice/PDX). Representative agarose gels of amplified products. Right numbers indicate amplicon size (in base pairs). Below numbers indicated the percentage of exon 8 exclusion. Mean ± SD.

## Discussion

Owing to their frequent deregulation in numerous cancers, targeting RNA Binding Proteins (RBPs) represents an opportunity in cancer treatment strategies. This last decade, splicing targeting drugs have emerged as a new class of promising anti-cancer drugs. However, their molecular mechanisms of action remain largely unknown. In addition, therapeutic strategies aiming at combine splicing targeting drugs with other therapies to enhance killing of tumor cells have been poorly explored. This might be particularly benefit in the context of tumor cells that acquire resistance to treatments and constitute the soil of relapse in patients. In this study, we identify SF3B1, a key component of the U2 snRNP component, as an Achilles’ heel of NSCLC cells with secondary resistance to platinum salts, a gold standard chemotherapy in lung cancer patients. Mechanistically, we demonstrated that inhibiting SF3B1 through pladienolide B treatment massively impacts the transcription and/or splicing of numerous genes involved in the DNA Damage Response, including *ATR*, *PRKDC* or *MLH3*, and counteracts the resistance to platinum salts both in NSCLC cells lines and PDXs. We also obtained “in vitro” evidence that combining pladienolide B with compounds targeting either RAD51, RAD52 or PARP-1/2, all key players of DNA repair pathways, could provide therapeutic benefits in NSCLC. As a whole, our data strengthen and extend to the lung cancer model recent studies published in other cancer types (60, 65–68) supporting the fact that SF3B1 inhibition or mutation massively and negatively impacts DNA Damage Response and repair and represents an actionable vulnerability to counteract resistance to current treatments. To our knowledge, our study also provides the first evidence of a growth inhibitory effect of pladienolide B, alone or in combination with platinum salts, in NSCLC PDXs.

We showed that pladienolide B-induced cell death in NSCLC cells with acquired resistance to platinum salts. Interestingly, it was previously shown that Small Cell Lung Carcinoma (SCLC) cells established from patients with prior chemotherapy or SCLC cells with acquired resistance to cisplatin display higher sensitivity to pladienolide B (69). Although the authors did not explore the molecular mechanisms involved in their study, these and our data strongly support a higher vulnerability to SF3B1 inhibition of lung cancer cells having received or escaping platinum-salts therapy. Despite pladienolide B derivatives, namely H3B-8800 and E7107, have demonstrated promising activities in various preclinical models supporting their enter in phase I/II clinical trials, their toxicity and insufficient efficacy have led to halt these clinical trials (35, 70, 71). These rather disappointing results underscore the necessity to identify patients who could better benefit from these therapies. It has been previously shown that these compounds preferentially kill spliceosome-mutant epithelial cells and hematological tumors that correlated with retention of short, GC-rich introns, enriched in genes encoding spliceosome components (28). SF3B1 mutations can also mediate the sensitivity to H3B-8800 in chronic lymphocytic leukemia (72). Additional determinants of the response to pladienolide B and derivatives have been identified (21). They include overexpression or hyperactivation of MYC which accelerates the synthesis of pre-mRNA, hence increasing the workload of the core spliceosome in its processing (16, 73), cohesin mutations in acute myeloid leukemia (AML) (68), or high levels in epithelial cells of anti-apoptotic members of the BCL-2 family, such as BCL-X_L_ or MCL-1 (74). In addition, elevated levels of SF3B1 protein increased susceptibility to pladienolide B in prostate and endometrial cancers (75, 76). In lung cancer, SF3B1 is rarely mutated. Nevertheless, we did DNA pyrosequencing to look for SF3B1(K700E) or U2AF1(S34F) mutation in our resistant H460 cells. We did not find any mutation (data not shown). We also did not evidence any variation of SF3B1 protein level between parental and resistant NSCLC cells (Fig 1e). Nevertheless, we noticed that H460 and A549 cells with acquired resistance to platinum salts displayed higher basal level of replicative stress compared to parental cells, as demonstrated by enhanced expression of P-ATR(Thr1989) protein or accumulation of RPA nuclear foci. We also provided evidence that pladienolide B activates an early transcription-dependent replicative stress response in these cells which was associated with enhanced number of RPA, P-53BP1(Ser1778) and γH2AX foci, as well as the accumulation of cells in G_2_/M. Interestingly, SF3B1^MUT^ leukemic cells display a defective response to PARP inhibitors-induced replication stress (77). These SF3B1^MUT^ cells down-regulate the cyclin-dependent kinase 2 interacting protein (CINP), leading to increased replication fork origin firing and loss of P-CHK1(Ser317) induction. This results in failure to resolve DNA replication intermediates and G_2_/M cell cycle arrest (77). In myelodysplatic syndromes, SF3B1 mutations also correlated with R-loop accumulation and DNA damage (78), although divergent results were recently published which demonstrated loss of R-loops with associated DNA replication stress in SF3B1-mutated MDS erythroblasts (79). We did not quantify R-loops in our cells. However, the negative effects of DRB on pladienolide B-induced P-DNA-PKcs(Ser2056), P-RPA32(Ser4/8) and γH2AX accumulation in NSCLC resistant cells suggested that enhanced transcription/replication conflicts could be involved in pladienolide B-mediated replicative stress. This remains to be further explored. As a whole, these and our results support the idea that cancer cells with enhanced susceptibility to replicative stress could be more vulnerable to SF3B1 inhibition. Among NSCLC PDXs, we did not find any common mutation(s) that could predict their differential response to pladienolide B (Table 1). However, as platinum salts-resistant NSCLC cells, we observed increased basal levels of P-ATR(Thr1989) protein in some PDXs responsive to pladienolide B alone or in combination. In addition, we noticed that the LCIM10 PDX which was the most responsive to pladienolide B was the only one with ATRX mutation. ATRX is implicated in DNA replication by assisting resolution of non-B DNA structures, such as G-quadruplex, thus protecting stalled replication forks from collapsing (80). In addition, ATRX deficiency increased sensitivity to agents which induce replication stress (81). Therefore, whether ATRX mutation could contribute to pladienolide B-induced tumor growth inhibition in NSCLC PDX remains to be further explored.

We demonstrated that pladienolide B enhances genomic instability in NSCLC resistant cells, indicating that these cells display DNA repair defects upon pladienolide B treatment. Consistently, after 24 hours exposure, pladienolide B shutdowns DNA damage signaling as evidenced by the decrease of both ATR and DNA-PKcs mRNA and/or protein levels. Importantly, we were able to confirm such a decrease of ATR and/or DNA-PKcs protein level upon pladienolide B treatment in two NSCLC PDXs. In these PDXs, this was not correlated with the decrease of *ATR* and/or *PRKDC* mRNA level (data not shown). We also showed that the effects of pladienolide B are associated with a strong impact on the expression, mostly down-regulation, and/or splicing, mostly exon skipping, of numerous DDR genes, including genes involved in all DNA repair pathways. Moreover, we showed that pladienolide B prevents the repair of I-SceI-induced double strand breaks (DSBs) by homologous recombination or c-NHEJ. These results are reminiscent with recent studies published in AML (68) or in Triple Negative Breast Cancer (TNBC) cells (60, 67), showing a widespread effect of pladienolide B on the expression and/or splicing of DNA damage repair genes. It has been proposed that these genes could be particularly sensitive to exon skipping given their long length and their large number of small exons per transcript which increases the number of exon-intron junctions thereby their dependency on efficient splicing (60). Importantly, we also showed that the impact on exon skipping of continuous exposure to pladienolide B is only transient. Most of the splicing events were detected after 6 hours treatment and returned to basal level after 24 hours. These data are also consistent with the results of Caggiano and colleagues who demonstrated that a short-pulse of pladienolide B leads to transient splicing defects but long-lasting depletion of DNA repair factors and cell death in TNBC cells (60). The authors speculated that this depletion could be linked to the elevated number of introns in DDR genes possibly supporting their incomplete or aberrant RNA processing during the recovery from splicing inhibition. Although we did not investigate this aspect in this study, we also observed that a 24 hours exposure to pladienolide B is sufficient to prevent NSCLC cell growth assessed after 10 days in clonogenic assay (data not shown). Nevertheless, among all the splicing events analyzed, skipping of *MLH3*-exon 8 and/or -exon 5 persisted until 48-72 hours treatment in resistant cells, and was also enhanced when pladienolide B was combined with cisplatin both in NSCLC and some NSCLC PDXs. Interestingly, *MLH3*-exon 8 skipping correlated with a decrease of MLH3 protein level in NSCLC cells, suggesting that *MLH3* transcripts devoid of exon 8 could be less stable and/or inefficiently translated. Although additional experiments are required to further deepen the role of MLH3 splicing in pladienolide B-induced cell death, it is important to mention that MLH3 is also part of a molecular signature that contains a common set of 275 transcripts differentially spliced and expressed upon SF3B1 downregulation in human AML (66).

One way to enhance the therapeutic efficiency of spliceosome inhibitors and to increase their tolerability is to use them in combination with additional therapies in order to exhibit synergistic cytotoxic effects. Consistently, pladienolide B demonstrated synergy with PARP inhibitors in TNBC (60, 67). E7107 and H3B-8800 also sensitized chronic lymphocytic leukemia (CLL) cells to the BCL-2 inhibitor venetoclax (72, 82). In this study, we found that pladienolide B restores sensitivity to platinum salts both in NSCLC cells with acquired resistance but also in some NSCLC PDXs that poorly respond to this chemotherapy. We also showed that combining pladienolide B with compounds targeting DDR, such as RAD51, RAD52 or PARP1/2 inhibitors, exhibits enhanced cell growth inhibitory effects as compared to each drug alone. Although these results remain to be confirmed “in vivo”, they unravel interesting therapeutic combinations that could provide therapeutic benefit in NSCLC. Together, our findings highly suggest that targeting SF3B1, alone or in combination with platinum salts or drugs targeting DNA repair pathways, could be exploited to counteract resistance in NSCLC patients.

## Declarations

### Ethics approval and consent to participate

Not applicable

### Consent for publication

Not applicable

### Availability of data and materials

The datasets used and analyzed during the current study are included in this published article and its supplementary information files or available from the corresponding author on reasonable request

### Authors’ contributions

B.E. designed the study, supervised the experiments and wrote the first draft of the manuscript. N.J.H and S.C. performed the majority of the *“in vitro”* experiments, analyzed and quantified the results with B.E., and prepared synthetic graphs. E.M and F.N performed all the *“in vivo”* experiments in PDXs. E.M. and D.D. designed the *“in vivo”* experiments, analyzed the results and prepared synthetic graphs. A.G. performed the SIRF experiments. H.P. and D.A. performed bioinformatic analyses of RNA-Seq. R.E.B. did the immunoblotting experiments in NSCLC PDXs. C.B. performed the high-throughput screening at CEA Grenoble. T.J. did correlative analyses based on CCLE database. B.E. and D.D. acquired fundings and edited the manuscript. All authors reviewed and approved the final version of the manuscript.

## Acknowledgements

We thank the platform of animal experiment of the Institut Curie. Part of this work has been performed at the CMBA platform - IRIG-DS-BGE-Gen&Chem-19 CMBA, CEA-Grenoble, F-38054 Grenoble (a member of GIS-IBISA and ChemBioFrance 20 National Research Infrastructure), which is supported by the LabEX GRAL (Grenoble Alliance 21 for Integrated Structural and Cell Biology), a program of the Chemistry Biology Health Graduate 22 School of Université Grenoble Alpes (ANR-17-EURE-0003). We thank the Microcell core facility of the Institute for Advanced Biosciences (UGA - Inserm U1209 - CNRS 5309), especially Jacques Mazzega, Solène Dufour and Mylène Pezet, for their assistance with microscopes/confocal microscope (J.M., M.P.) and flow cytometer (S.D.), as well as M.P. and S.D. for their help in quantifying γH2AX nuclear foci and SIRF signals using specific macro and tools. This facility belongs to the IBISA-ISdV platform, member of the national infrastructure France-BioImaging supported by the French National Research Agency (ANR-10-INBS-04).

## Supplementary Figures Legends

**Supplementary Figure 1. Cytotoxic effects of pladienolide B in various NSCLC cell lines including cellular models with acquired resistance to EGFR-TKIs.** (a) PrestoBlue cell viability assay in indicated NSCLC cell lines treated with increasing concentrations of pladienolide B (nM) for 72 hours. (b) MTS assay in H460S/R or A549 S/R cells treated or not for 96 hours with increasing concentrations of cisplatin (µM). Mean ± SD. n=3. (c) PrestoBlue cell viability assay in NSCLC cell lines with acquired resistance to EGFR-TKIs and treated with increasing concentrations of pladienolide B (nM) for 72 hours. HCC827-DACOR correspond to HCC827-EGFR mutated cells with acquired resistance to dacomitinb. PC9-GEFR, PC9-DACOR, and PC9-OSIR represent PC9-EGFR mutated cells with acquired resistance to gefitinib, dacomitinib or osimertinib, three distinct generation of EGFR-TKIs, respectively. Mean ± SD.

**Supplementary Figure 2. Effects of pladienolide B on tumor growth of NSCLC Patient-Derived-Xenografts (PDXs).** (a) Absence of toxicity in mice following pladienolide B treatment alone or in combination with cisplatin. Mice (n = 3) were treated intra-peritoneally with 7 mg/kg pladienolide B at days 1 and 3 followed by 5 mg/kg at days 5 and 8 (left panel), or mice (n = 4) received 5 mg/kg pladienolide B at days 3, 5, 8 or 10 and 4 mg/kg cisplatin intra-peritoneally 3 times/week between day 1 to 21 (right panel). Mice were weighted every two days. (b) Overall Rate Response (ORR) was calculated in different PDXs receiving either vehicle (control), 2.5 mg/kg or 5 mg/kg pladienolide B intra-peritoneally at days 1, 4, 8 and 11. Two-tails Mann-Whitney t test. * p< 0.05. (c) Relative Tumor Volume (RTV) was calculated at different times in mice engrafted with 8 distinct NSCLC PDXs (n = 3 to 7 mice/PDX) and treated as in (b). Mean ± SEM. (d) Probability of tumor progression (RTV = 4) in 7 PDXs (excepted LCIM1) treated as in (b, c). Log-rank t test. ** p< 0.01.

**Supplementary Figure 3. Pladienolide B down-regulates the expression of genes involved in DNA metabolic process.** (a, b) H460S (a) or H460R (b) cells were treated for 8 hours with 3 nM pladienolide. GO-term enrichment analysis using Metascape of Differentially Expressed Genes (DEGs) in treated condition as compared to control. DEGs either down- or up-regulated in H460S or H460R cells according to absolute Log2 FC ≥ 1.

**Supplementary Figure 4. Pladienolide B regulates the splicing (exon skipping/mutually exclusive exon) of genes involved in DNA Damage Response in NSCLC cells.** GO-term enrichment analysis using Metascape of Differentially Spliced Genes (DSGs) with exon skipping or mutually exclusive exons in H460S (a) or H460R (b) cells. p value ≤ 0.05 and absolute deltaPSI ≥ 20%.

**Supplementary Figure 5. Pladienolide B regulates expression and splicing of genes involved in DNA Damage Response in parental and resistant H460 cells.** (a) Upper panel: specific and common DSGs (p value ≤ 0.05, absolute deltaPSI ≥ 20%) and DEGs (p value ≤ 0.05, absolute log2FC ≥ 0.4) in H460S treated with 3 nM pladienolide B for 8 hours as compared to untreated cells. Lower panel: GO-term enrichment analysis using Metascape of common DSGs and DEGs. (b) Upper panel: specific and common DSGs (p value ≤ 0.05, absolute deltaPSI ≥ 20%) and DEGs (p value ≤ 0.05, absolute log2FC ≥ 0.4) in H460R treated with 3 nM pladienolide B for 8 hours as compared to untreated cells. Lower panel: GO-term enrichment analysis using Metascape of common DSGs and DEGs.

**Supplementary Figure 6. Pladienolide B induces exon skipping of genes involved in DNA repair.** RT-PCR analyses of indicated exon skipping events. Representative agarose gels of amplified products. Right numbers indicate amplicon size (in base pairs).

**Supplementary Figure 7.** Mice were engrafted with 7 distinct NSCLC PDXs and received either vehicle (control), cisplatin (4 mg/kg, ip, 1 x/3 weeks), pladienolide B (5 mg/kg, ip, D1-D4-D8-D11) or a combination of pladienolide B + cisplatin. (a) Probability of tumor progression (RTV=4). Log-rank t test. * p< 0.05, ** p< 0.01, *** p< 0.001. (b) Overall Rate Response (ORR) of 7 NSCLC PDXs treated in the same conditions as in (a). (c) Relative Tumor Volume (RTV) was calculated at different times in mice treated in the same conditions as in (a) (n = 3-4 mice/each PDX). Mean ± SEM. Two-tail Mann-Whitney t test. (d) Immunoblotting of P-ATR(Thr1989) protein in various NSCLC PDXs including LCF2, LCIM13, LCF15, LCF26, LCIM1, ML5LC66, ML1LC2, and LCIM10. GAPDH was used as a loading control.

**Supplementary Table S1.** List of all antibodies used in this study.

**Supplementary Table S2.** List of all primers used in RT-PCR validation.

