## Supplementary figures and images for "The SF3B1 inhibitor pladienolide B massively inhibits DNA damage signaling and repair and counteracts resistance to platinum salts in Non-Small Cell Lung Cancer"

### Supplementary Figure 1

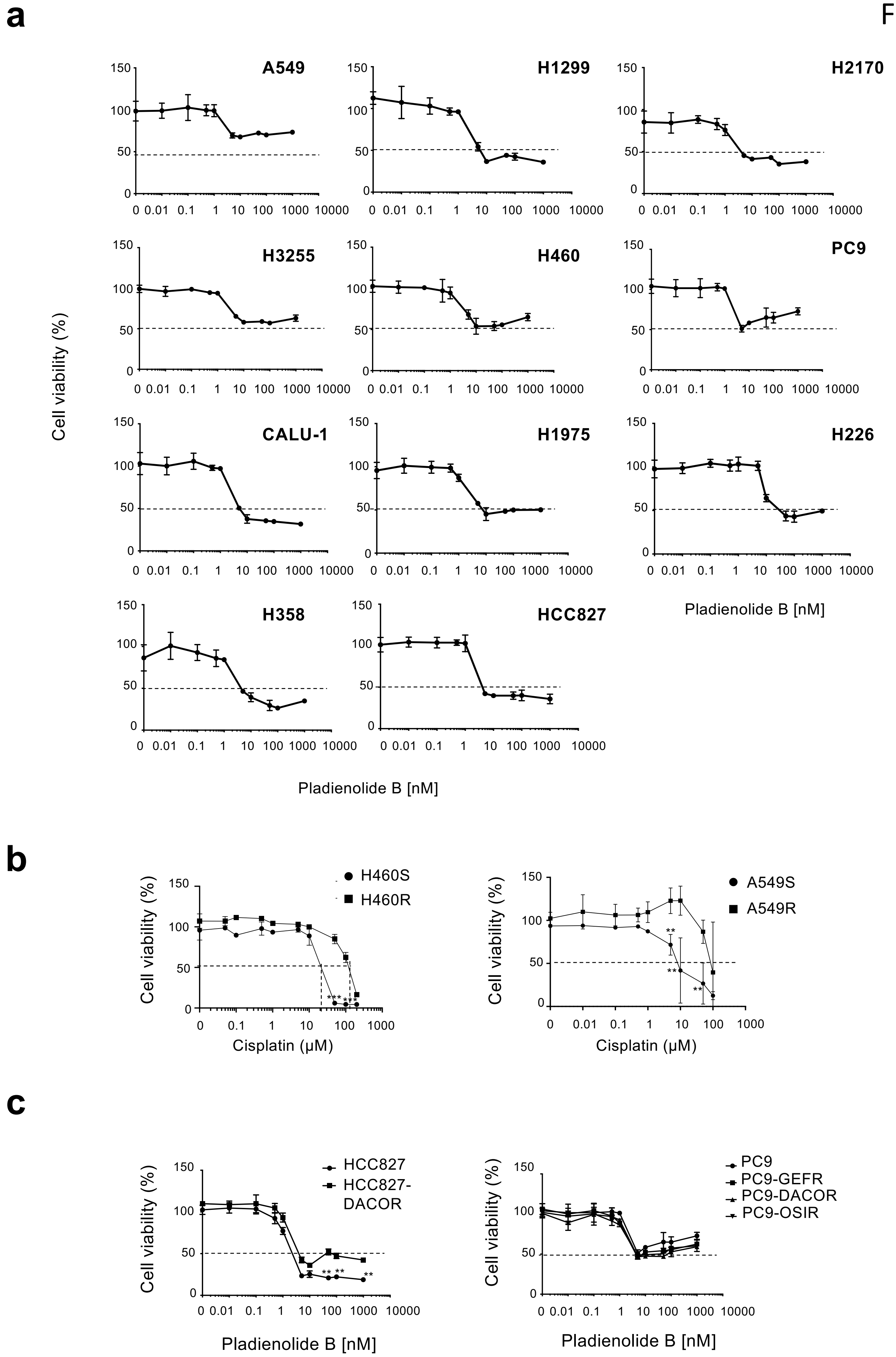

### Supplementary Figure 2

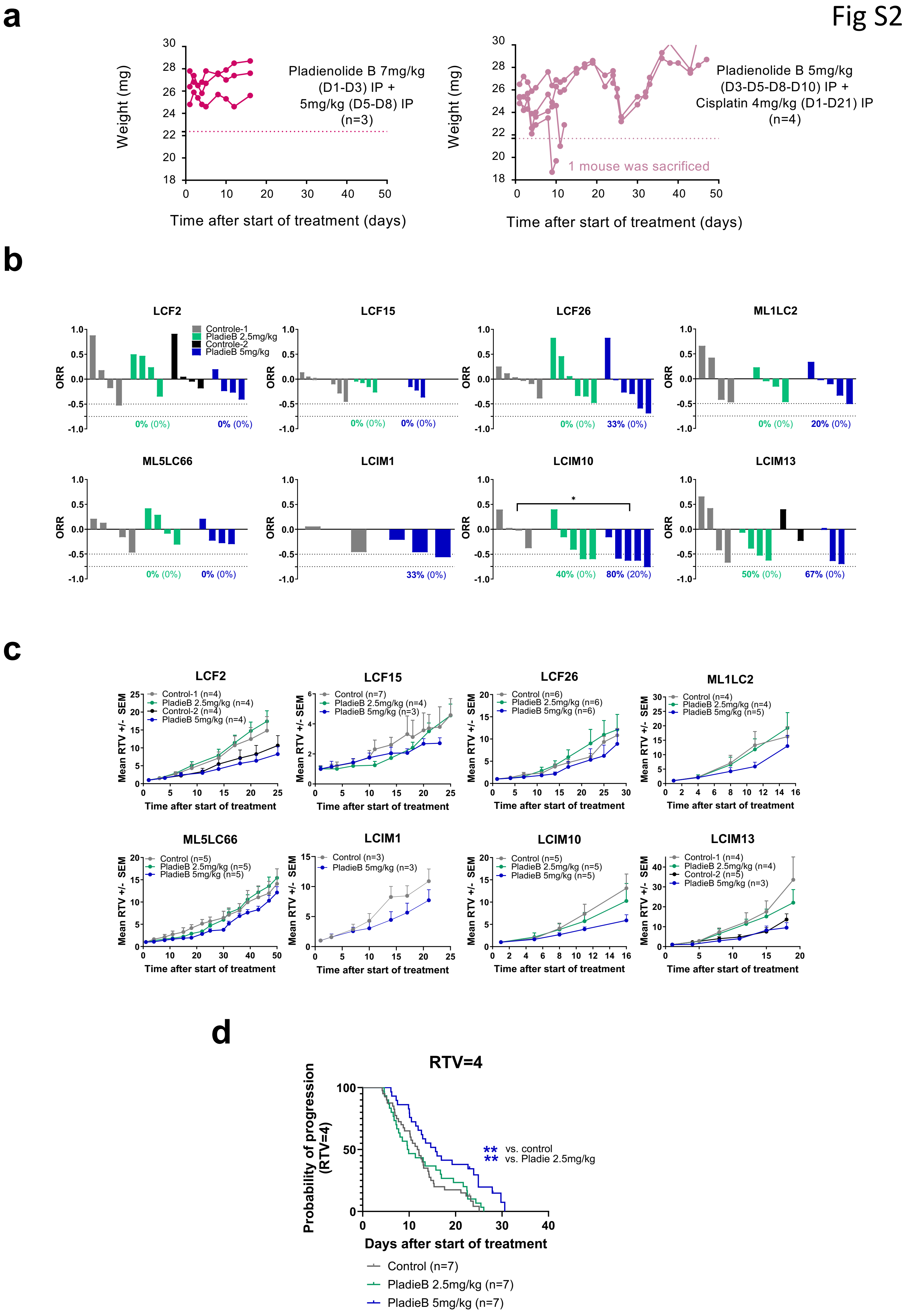

### Supplementary Figure 3

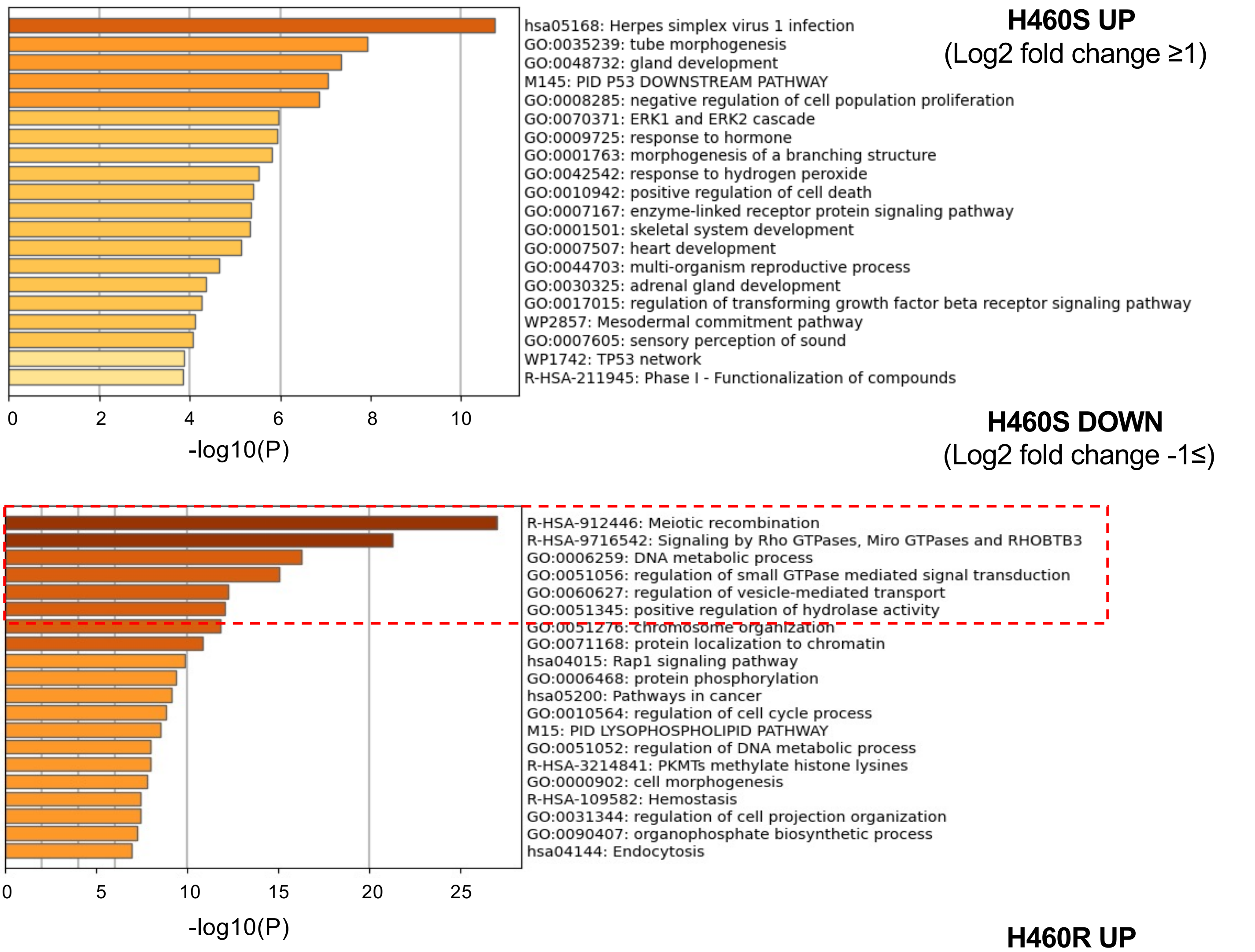

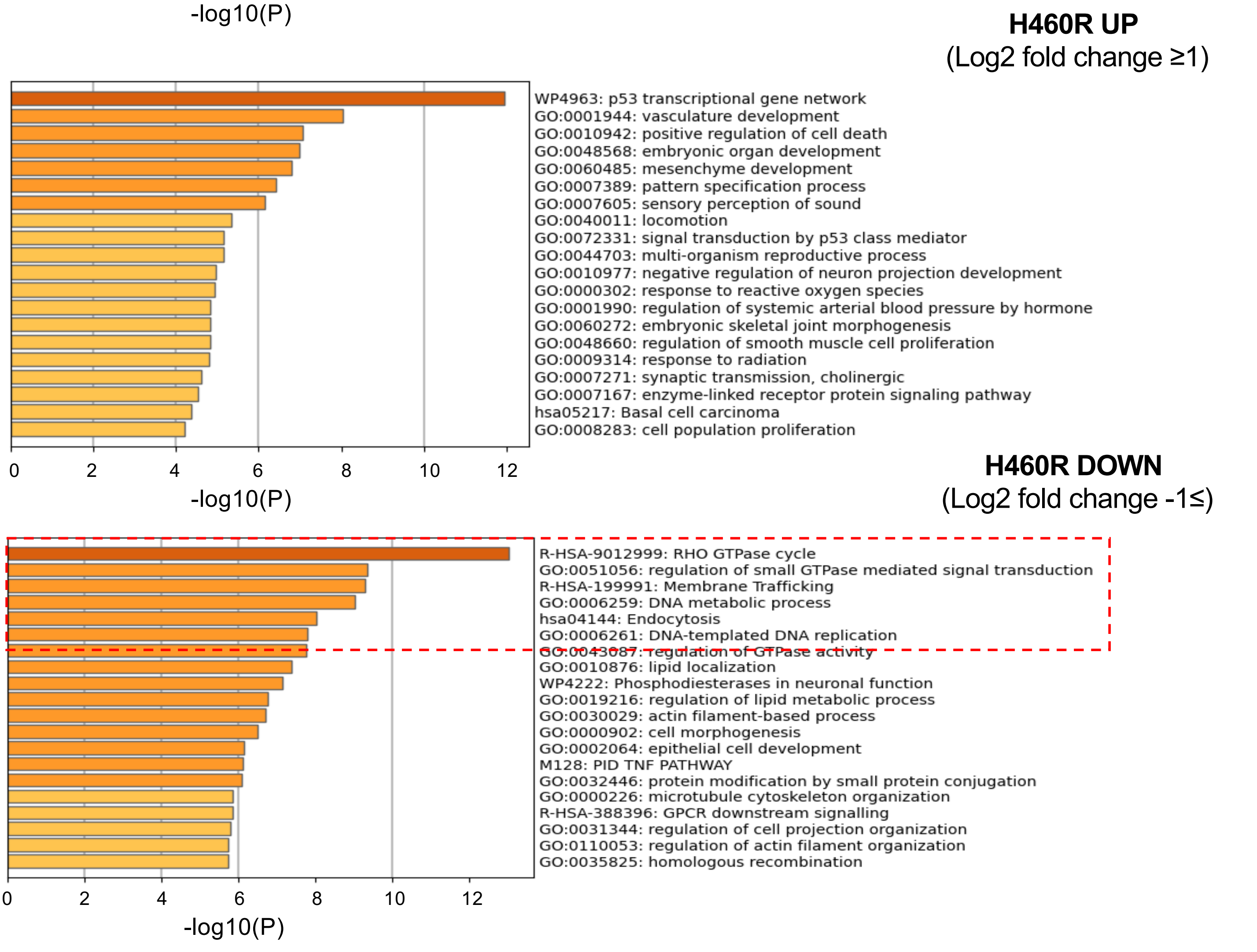

### Supplementary Figure 4

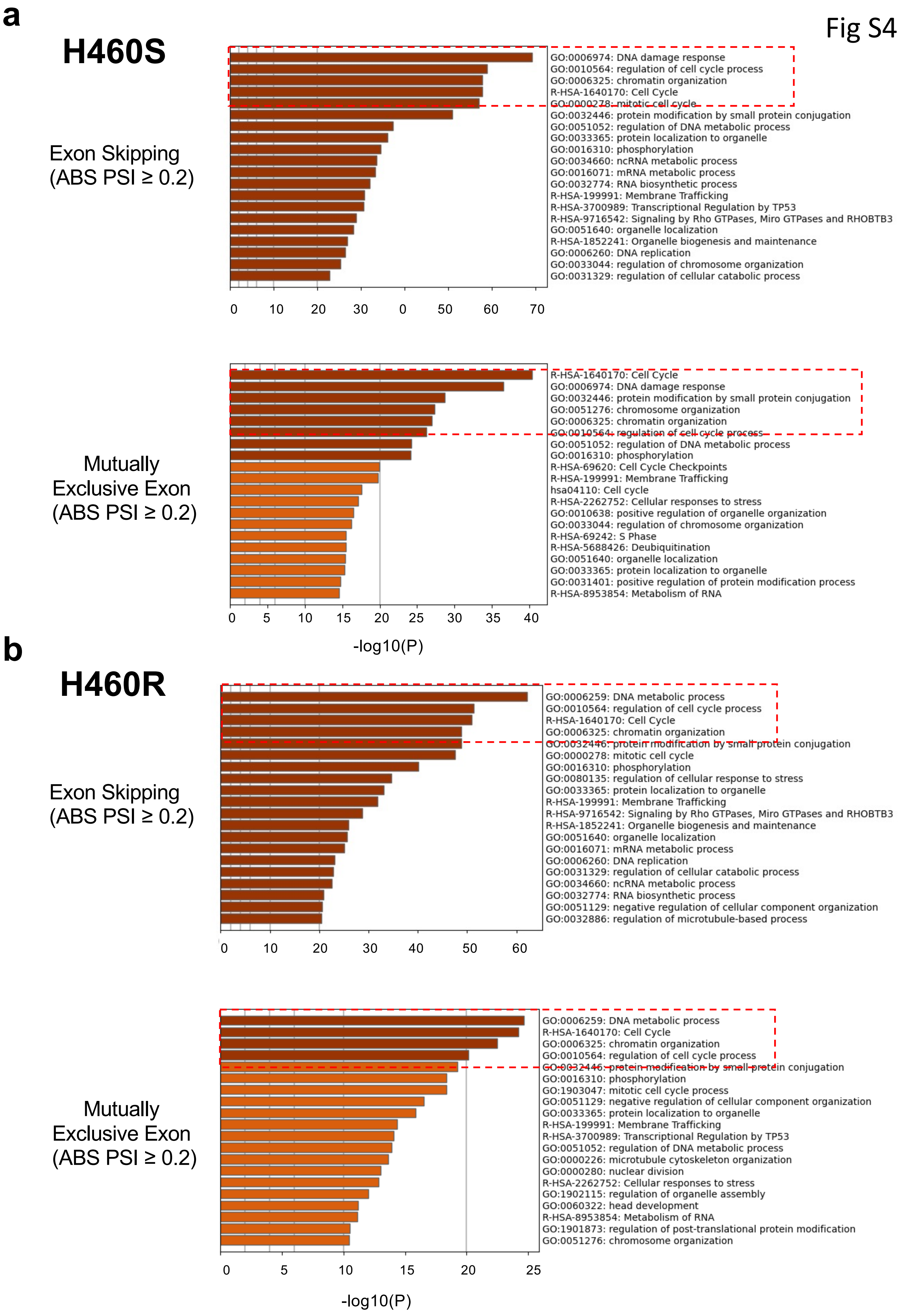

### Supplementary Figure 5

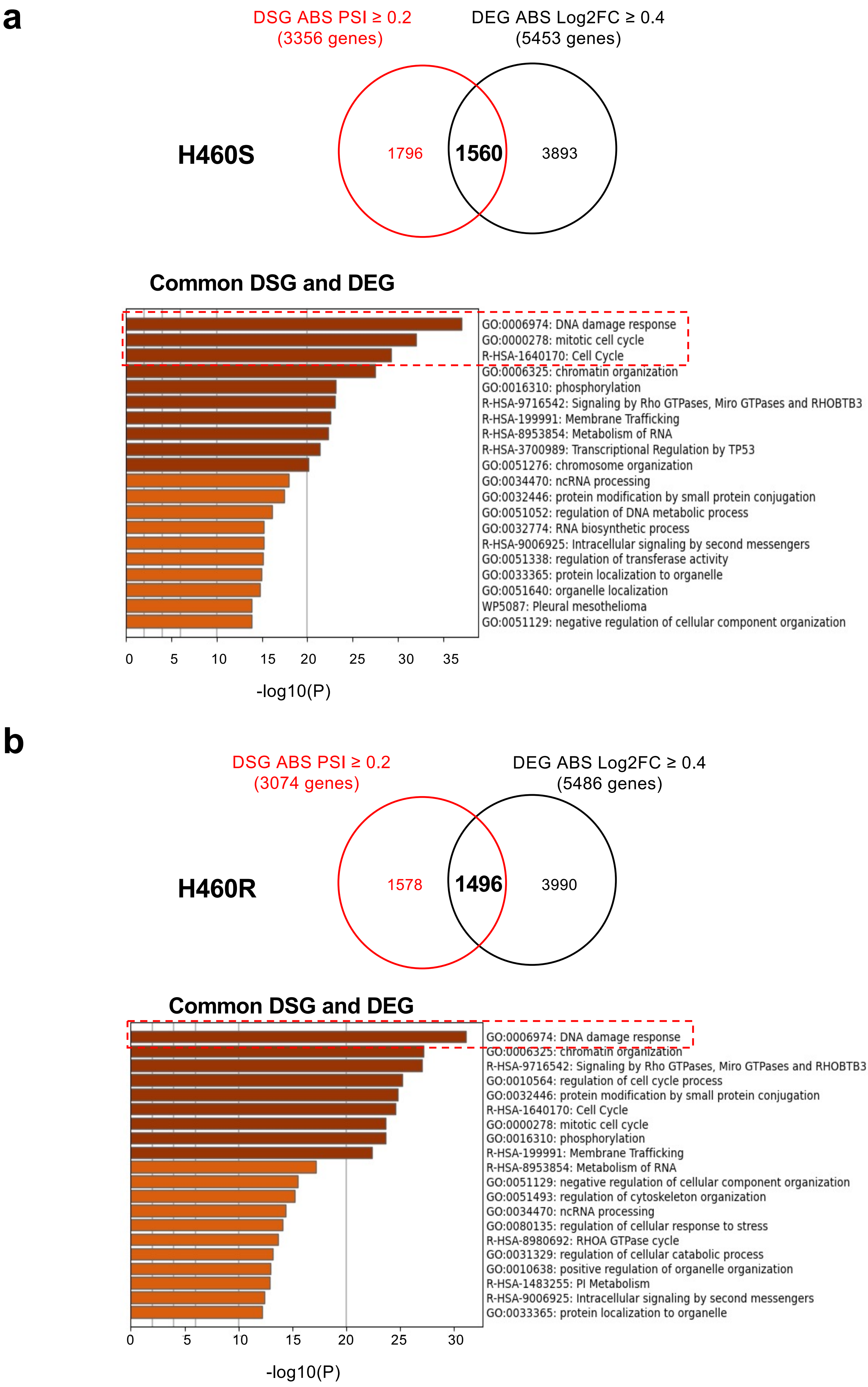

### Supplementary Figure 6

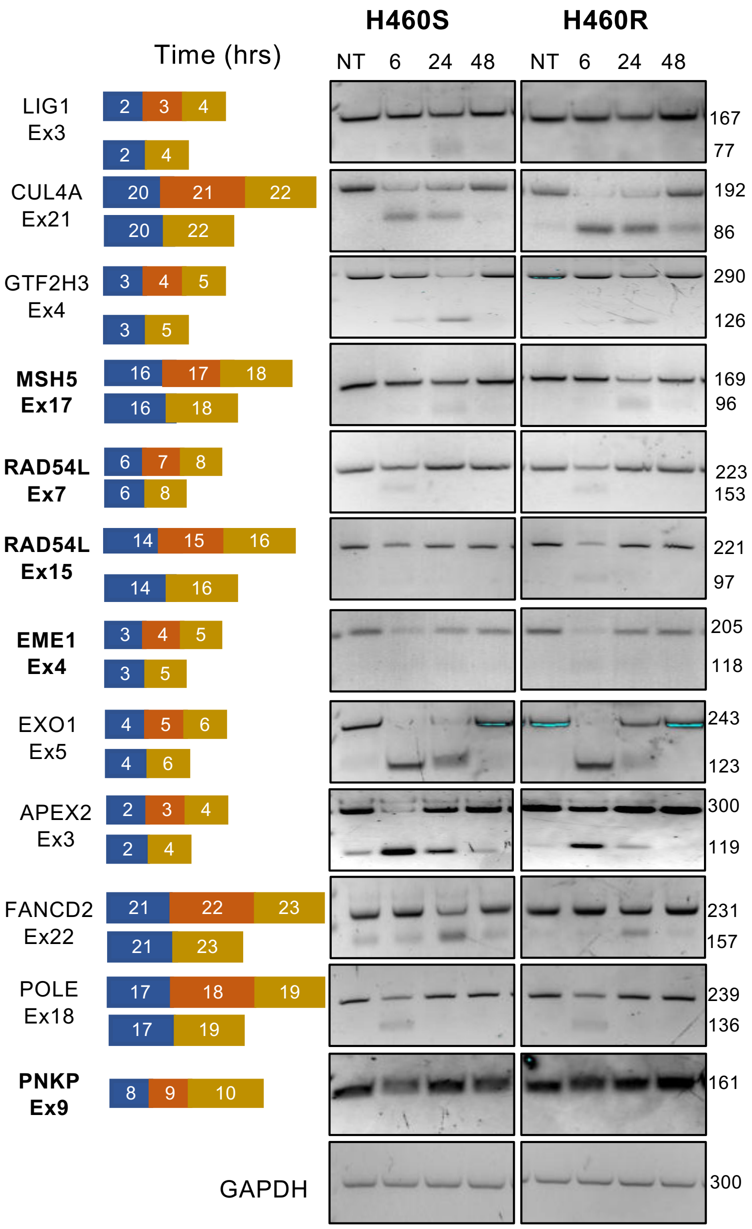

### Supplementary Figure 7

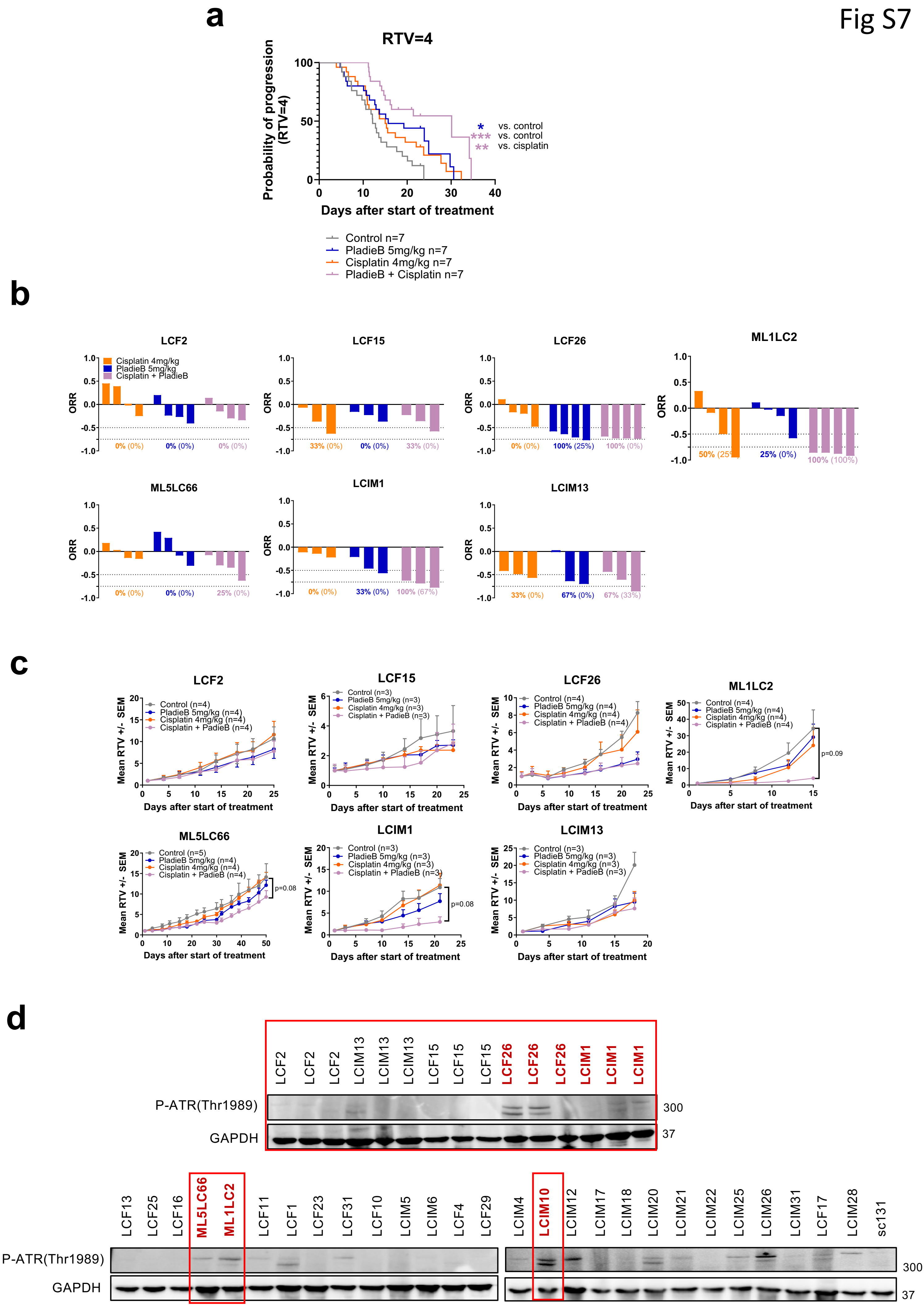
