## Supplementary Table 1 for "The SF3B1 inhibitor pladienolide B massively inhibits DNA damage signaling and repair and counteracts resistance to platinum salts in Non-Small Cell Lung Cancer"

**Supplementary Table S1.** List of antibodies used in this study.

| Antibody | Source | Reference |
|---|---|---|
| CLEAVED CASPASE-3 | Cell Signaling Technologies | Cat#9661 |
| ATR | Cell Signaling Technologies | Cat#2790 |
| CD4 | BD biosciences | Cat#553046 |
| DNA-PKcs | Santa Cruz Biotechnologies | Cat#sc-390849 |
| GAPDH | Santa Cruz Biotechnologies | Cat#sc-47724 |
| HA | Cell Signaling Technologies | Cat#2367 |
| KU80 | Cell Signaling Technologies | Cat#2180 |
| MLH3 | Santa Cruz Biotechnologies | Cat#sc-25313 |
| PCNA | Santa Cruz Biotechnologies | Cat#sc-56 |
| P-53BP1(Ser1778) | Cell Signaling Technologies | Cat#sc-25313 |
| P-ATR(Thr1989) | Cell Signaling Technologies | Cat#30632 |
| P-CHK1(Ser345) | Cell Signaling Technologies | Cat#2348 |
| P-DNA-PKcs(Ser2056) | Cell Signaling Technologies | Cat#68716 |
| P-H2AX(Ser139) | Cell Signaling Technologies | Cat#2577 |
| P-H2AX(Ser139) | Millipore | Cat#05-636 |
| P-RPA32(S4/S8) | Bethyl Laboratories | Cat#A300-245A |
| RPA32 | Santa Cruz Biotechnologies | Cat#52448 |
| SF3B1 | Proteintech | Cat#276841AP |
| α-TUBULIN | Santa Cruz Biotechnologies | Cat#sc-23948 |

WB: western blot
IF : immunofluorescence
FC: Flow cytometry
SIRF: in situ analysis of proteins at nascent replication forks

| Application |
| --- |
| WB |
| WB |
| FC |
| WB |
| WB |
| WB |
| WB |
| WB |
| WB, SIRF |
| WB |
| IF |
| WB |
| WB |
| WB |
| IF |
| WB |
| WB |
| WB |
| WB |
