## Supplementary Table 2 for "The SF3B1 inhibitor pladienolide B massively inhibits DNA damage signaling and repair and counteracts resistance to platinum salts in Non-Small Cell Lung Cancer"

| PCR Primers | | | |
|---|---|---|---|
| **Gene** | **Primer** | **Exon** | **Sequence (5'->3')** |
| APEX2 | Forward | Exon 2 | CCCCTGGCTATCGTTGAGG |
| (Exon 3) | Reverse | Exon 4 | GGGGCAGTACACGTTGATTA |
| CDK7 | Forward | Exon 2 | CCACCGTTTACAAGGCCAGA |
| (Exon 4) | Reverse | Exon 5 | TATTTGGATGACTTAGCTCCTGT |
| CUL4A | Forward | Exon 20 | GGTCGAAAACTTCAGTGGCA |
| (Exon 21) | Reverse | Exon 22 | CGTTCTGCGCAATTCACTATC |
| GTF2H4 | Forward | Exon 8 | CAGCTTCTCTACTCTGGGCA |
| (Exon 9) | Reverse | Exon 10 | ACGACAATGAAACCTGGCTG |
| GTF2H3 | Forward | Exon 3 | CGTGATGGTGCTGGGAAATT |
| (Exon 4) | Reverse | Exon 5 | GGGATCCTGCCAGCAAAGT |
| DCLRE1C | Forward | Exon 7 | TGAAATCGAGACTCCTACCCAG |
| (Exon 8) | Reverse | Exon 9 | CTCCTTGCGCCAATCTGAAG |
| ATR | Forward | Exon 29 | ACATGAAAGCCTTGGCTTGC |
| (Exon 30) | Reverse | Exon 31 | TGCTTCCACTCTGTACGTGT |
| CHEK1 | Forward | Exon 3 | GGAGAAGGTGCCTATGGAGA |
| (Exon 4) | Reverse | Exon 5 | GTTGATGGAAGAATCTCTGAGCA |
| CHEK1 | Forward | Exon 8 | GACCAACCCAGTGACAGCT |
| (Exon 9) | Reverse | Exon 10 | GACTCTGACACACCACCTGA |
| EME1 | Forward | Exon 3 | CTAGGAGCACTGCAGACCAT |
| (Exon 4) | Reverse | Exon 5 | TCATAGTGCTGTCCAGGCTT |
| EXO1 | Forward | Exon 2 | GGATATTTGCCTGGCCCAG |
| (Exon 3) | Reverse | Exon 4 | TGTATCCCCATGGTGCCAAA |
| EXO1 | Forward | Exon 4 | GGATACATATTGCTGGCTTCACA |
| (Exon 5) | Reverse | Exon 6 | AGACTCCAGGCCACTAACAA |
| FANCD2 | Forward | Exon 21 | GACCTCCTTGTTGCAGTTGG |
| (Exon 22) | Reverse | Exon 23 | TCCAGTCCGTACAGTGCTTT |
| LIG1 | Forward | Exon 2 | AGGGAGAATTCTGACGCCAA |
| (Exon 3) | Reverse | Exon 4 | CTCTCGGACACCACTCCATT |
| MLH3 | Forward | Exon 4 | GACAATCCAGTATTTGCCCGT |
| (Exon 5) | Reverse | Exon 6 | CCTCGCCATTCTCTTCAGTC |
| MLH3 | Forward | Exon 6/7 | GACTGAAGAGAATGGCGAGG |
| (Exon 8) | Reverse | Exon 9 | TTTTCCGACCAGAGCCTTGT |
| MSH5 | Forward | Exon 16 | TCCAGCTCTTTCGGGACATT |
| (Exon 17) | Reverse | Exon 18 | AGTCCCATCAGTCTTCGCTT |
| PARP2 | Forward | Exon 5 | GTGTTTGGATGAGATGGGGC |
| (Exon 6) | Reverse | Exon 7 | TCCAGGCACCTTCTCAAACT |
| PARP2 | Forward | Exon 14 | TGCCAATTACTGCTTTGCCT |
| (Exon 15) | Reverse | Exon 16 | CACTTGCTGGTCCTAATGGC |
| PNKP | Forward | Exon 8 | CACGCACGCAGGCTTGTA |
| (Exon 9) | Reverse | Exon 10 | AGTCTTTCTTCTTCCGCCCC |
| POLD3 | Forward | Exon 6 | GTTTCCCAGCAGCCCAAAG |
| (Exon 7) | Reverse | Exon 8 | TCACTGCTTGTTCTGAGTCCA |
| POLE | Forward | Exon 3 | CTTAGGCAGTGCAGTGGATT |
| (Exon 4) | Reverse | Exon 5 | TTGCCCTGAAACTTCTTGGAG |
| POLE2 | Forward | Exon 10 | ACCCACTGAGCCCTCTAGTA |
| (Exon 11) | Reverse | Exon 12 | GGTTGGAGGTGCTGGTGAAT |
| POLE | Forward | Exon 17 | ACCCCAACATCATCCTGACC |
| (Exon 18) | Reverse | Exon 19 | CAGTTCATGAAAGGCCCGAG |
| POLE3 | Forward | Exon 2 | ACCAGGATCATCAAGGAGGC |
| (Exon 3) | Reverse | Exon 4 | GTAACGAACCGCTGGAACTC |
| RAD52 | Forward | Exon 11 | CCGTCAGCTGAGAAGAGTGA |
| (Exon 12) | Reverse | Exon 13 | GGAGTCACAGCCCACTTTTC |
| RAD54L | Forward | Exon 6 | TGCCTTGGTTCTGTATGAGC |
| (Exon 7) | Reverse | Exon 8 | CGTCTTTCCTAGGCCCATCT |
| RAD54L | Forward | Exon 14 | TGGTTACAGCTCTAAGGCCC |
| (Exon 15) | Reverse | Exon 16 | CTACAACCTTGGCTCGCTTC |
| REV1 | Forward | Exon 5 | GCCACAAATCTTCCCAATGC |
| (Exon 6) | Reverse | Exon 7 | TCCAACTGTTCATGCCATTGA |
| REV1 | Forward | Exon 7 | GGTCTGCACTTGTTGTAACTGA |
| (Exon 8) | Reverse | Exon 9 | CCCTTCCTGTGCCTCTGTTA |
| REV3L | Forward | Exon 18 | GGGCAAGAACTGGGAAATGT |
| (Exon 19) | Reverse | Exon 20 | CTGTGGGGTATTTACTGCTGC |
| REV3L | Forward | Exon 32 | TGGTCAGGAAATTGCCGAAG |
| (Exon 33) | Reverse | Exon 34 | TCTCTCGTTTCAAATAGCAGCT |
| WRN (Exon | Forward | Exon 13 | CAGCACCCAATGAAGAGCAA |
| 14) | Reverse | Exon 15 | GGGGAGAGATAACAAGGCCA |
